# CSI-SSU: Phylogenetic contamination screening of genomic datasets, demonstrated on the Protist 10,000 Genomes (P10K) database

**DOI:** 10.1101/2025.08.19.670927

**Authors:** Alfredo L. Porfirio-Sousa, Robert E. Jones, Matthew W. Brown, Daniel J. G. Lahr, Alexander K. Tice

## Abstract

**Background:** Genomic data are essential for uncovering the evolutionary history, ecological roles, and diversity of life. Yet, diverse microbial eukaryotes, predominantly unicellular and traditionally referred to as protists, remain critically underrepresented in genomic repositories, limiting our ability to address fundamental questions in eukaryotic evolution. The Protist 10,000 Genomes (P10K) initiative seeks to fill this gap by generating and compiling genomic and transcriptomic data for a wide range of microbial eukaryotes. However, large-scale sequencing efforts face persistent challenges, including contamination and imprecise taxonomic identification, particularly for poorly studied taxa that require specialized taxonomic expertise. To ensure the reliability of these resources, robust and scalable approaches for taxonomic identification and contamination screening are essential.

**Results:** We developed CSI-SSU (https://github.com/AlexTiceLab/CSI-SSU), a command-line tool for Contaminant Sequence Investigation (CSI) that uses small subunit ribosomal RNA (SSU) sequences, chimeric sequence detection, and phylogenetic placement to rapidly identify, retrieve, and classify SSU sequences from eukaryotic genomic-level assemblies. CSI-SSU incorporates a curated SSU reference dataset representing the major known eukaryotic supergroups, with sequences and taxonomic nomenclature derived from the Protist Ribosomal Reference (PR2) database. In addition to detecting contaminant sequences, CSI-SSU enables approximate taxonomic assignment of the target lineage in each assembly, with resolution constrained by the current diversity represented in PR2. To further assess potential bacterial contamination, CSI-SSU employs bacterial BUSCO searches as a proxy. We demonstrate CSI-SSU utility and performance by screening 2,960 genomic-level assemblies spanning a broad diversity of eukaryotes from P10K. CSI-SSU efficiently detected non-target eukaryotic SSU sequences, revealing cross-group contamination. Classifications also corroborated or refined the original taxonomic assignments, with resolution depending on PR2 representation. Bacterial BUSCO searches indicated bacterial contamination. Independent SSU and COI phylogenies of Amoebozoa supported CSI-SSU classifications, highlighting its accuracy and sensitivity.

**Conclusion:** CSI-SSU provides a scalable and reproducible framework for phylogenetically informed contamination screening and taxonomic validation of genomic and transcriptomic data. Coupling phylogenetic placement with contamination detection enabled us to distinguish high-quality P10K datasets from those requiring decontamination or additional sequencing before downstream use. These findings serve as a reference for future analyses and guide further sequencing efforts to expand the taxonomic diversity of microbial eukaryotes at the genomic level. Addressing imprecise taxonomic assignments, contamination, and reproducibility in genomic-level datasets will enhance the value of these resources and facilitate studies illuminating the evolution and diversification of eukaryotic life.

## Introduction

Genomic-level data are essential for advancing our understanding of the evolution of life on Earth [1, 2]. High-quality genome and transcriptome sequences enable comparative analyses that reveal patterns of genomic evolution, including gene family expansions, horizontal gene transfers, and changes in genome organization and regulatory systems [3, 4]. Such data also clarify phylogenetic relationships through multi-gene and genome-scale reconstructions [5, 6]. Moreover, genomic analyses uncover ecological interactions by identifying metabolic pathways, symbiotic associations, and genetic adaptations to specific environmental conditions [7]. Consequently, genomics has become central to studying life’s diversity and complexity.

While genomics has progressed rapidly for major eukaryotic groups such as plants, animals, fungi, and other traditional model systems, most microbial eukaryotes (commonly referred to as protists) remain vastly underrepresented [1, 2]. Although they comprise most of eukaryotic diversity, the majority of lineages within this highly diverse paraphyletic assemblage still lack genomic data [2, 5]. Over the past decade, several initiatives have contributed to fill this gap, such as the Marine Microbial Eukaryote Transcriptome Sequencing Project (MMETSP) for marine microbial genomics [8], the Tree of Life Programme for eukaryotic genome sequencing [9], and the One Thousand Plant Transcriptomes initiative (1KP) for plants, including single-celled algae [10]. Similarly, the Protist 10,000 Genomes (P10K) initiative aims to address the underrepresentation of microbial eukaryotes in genomic databases by generating new genomic data (i.e., genomes and transcriptomes) and compiling previously available data from a wide array of lineages, coupled with taxonomic identification and decontamination procedures [2]. This large-scale effort, consolidated in the P10K database, represents an unprecedented genomic resource for protists and potentially provides a valuable foundation for achieving a comprehensive and integrative view of eukaryotic evolution.

However, to fully exploit the potential of this newly available genomic resource for protists, three key challenges must be considered: (1) accurate taxonomic identification, (2) the presence of contaminated genomic data resulting from non-target eukaryotic contamination, and (3) data interpretation and reproducibility [11, 12]. Specifically, the taxonomic identification of the P10K database relies on small subunit ribosomal DNA (SSU rDNA) retrieval and BLAST similarity searches against curated SSU databases, including SILVA, PR^2^, and NCBI’s NT (nucleotide sequence) database [2]. Although informative, a BLAST-based approach is less accurate than phylogeny-based methods for taxonomic identification and may lead to a less precise classification of the target groups. Regarding contamination, protists typically inhabit environments with high eukaryotic microbial diversity, several prey on other organisms, many host endosymbionts, and most are often difficult to isolate as monoeukaryotic cultures or single cells [13–15]. As a result, genomic-level sequencing efforts might produce contaminated assemblies that contain not only the genome of the target organism but also sequences from other eukaryotic organisms [16, 17]. While P10K applies a decontamination strategy to remove bacterial, archaeal, viral, fungal, and other eukaryotic contaminants from ciliate data, for other protist groups only bacterial, archaeal, and viral contaminants are filtered [2]. Finally, the P10K database currently lacks photo-documentation and detailed information, for instance, about the sequencing platforms used for each sample (i.e., Illumina, Oxford Nanopore, or PacBio Sequel II), which are essential for deeper interpretation and reproducibility of the data. In this context, accurate taxonomic identification, assessment of eukaryotic contamination, and the availability of more detailed sample information are most essential for the effective downstream use of data in the P10K database.

Here, we introduce CSI-SSU, a command-line tool for Contaminant Sequence Investigation (CSI) that uses small subunit ribosomal RNA (SSU) sequences, chimeric sequence detection via VSEARCH, bacterial BUSCO searches, and phylogenetic placement via pplacer to rapidly identify, classify, and screen eukaryotic genomic resources. We applied CSI-SSU to the Protist 10,000 Genomes (P10K) database to evaluate taxonomic identification and contamination across diverse eukaryotic lineages. Incorporating a curated SSU reference dataset representing the major known eukaryotic supergroups, CSI-SSU detects non-target and chimeric sequences, enables approximate taxonomic assignment of assemblies, and assesses potential bacterial contamination using BUSCO searches. Phylogenetic analyses focused on Amoebozoa as a case study, using SSU and cytochrome c oxidase subunit I (COI) markers, validated CSI-SSU classifications and identification of high-quality assemblies from those requiring decontamination or additional sequencing. This framework provides a scalable, reproducible approach for improving the reliability, usability, and interpretability of large genomic datasets, offering a proof-of-concept for phylogenetically informed contamination screening and taxonomic validation across all major eukaryotic lineages using CSI-SSU tool (https://github.com/AlexTiceLab/CSI-SSU).

## Methods

### CSI-SSU tool

The full mode of the CSI-SSU tool performs automated SSU (18S) screening on genomic-level data using BLAST v2.17.0, MAFFT v7.526, pplacer v1.1.alpha20, and VSEARCH v2.30.5 [18–21] (**Figure S1**). CSI-SSU requires a single input: a nucleotide assembly (e.g., genome, transcriptome, metagenome, or metatranscriptome) to be screened. To identify and retrieve SSU sequences from target assemblies, CSI-SSU implements a similarity search using BLAST+ v2.17.0, based on a curated reference SSU query dataset. Query sequences were selected from the Protist Ribosomal Reference (PR2) database to capture the broad diversity of major eukaryotic lineages (https://github.com/AlexTiceLab/CSI-SSU/tree/main/csi_ssu/data/queries). For each input assembly, CSI-SSU first constructs a local BLAST database using the *makeblastdb* command. The reference SSU queries are then used in a *blastn* search to identify candidate SSU sequences within the assembly. BLAST results are subsequently parsed, and SSU sequences are retrieved using the Python script *parse_blast*.*py*, which extracts full subject sequences and corrects their orientation based on BLAST coordinates. Once retrieved, SSU sequences are subjected to phylogenetic placement.

To perform phylogenetic placement, CSI-SSU provides curated reference alignments and phylogenetic trees for nine major eukaryotic supergroups (Amoebozoa, Excavata, TSAR, Archaeplastida, Cryptista, Haptista, CRuMs, Provora, and Obazoa), derived from the PR2 database. To construct these reference datasets, we downloaded the PR2 SSU database (version 5.1.0) on September 9th, 2025. Redundancy was reduced using CD-HIT v4.8.1 with the parameter “-c 0.95,” corresponding to a global similarity threshold of 95%. For each target supergroup, the reference dataset included: (i) all sequences assigned to the focal supergroup in PR^2^, (ii) sequences labeled as Eukaryota_X, and (iii) five representative sequences from each class of all other supergroups available in PR^2^. Taxonomic assignments followed the PR2 classification as accessed on September 9th, 2025. For each supergroup-specific FASTA file, sequences were aligned using MAFFT *(--auto*) and trimmed with trimAl v1.2 using a gap threshold of 0.3 (*trimal -gt 0*.*3*). Preliminary phylogenetic trees were inferred using RAxML with 100 bootstrap replicates. These trees were manually inspected to remove putative contaminants and long-branch sequences. The untrimmed, curated datasets were then realigned using MAFFT (*--auto*) and trimmed again with trimAl (*-gt 0*.*3*). The resulting curated alignments and phylogenetic trees constitute the reference datasets used for SSU sequence placement.

CSI-SSU performs phylogenetic placement of retrieved SSU sequences using pplacer. Retrieved sequences are added to the selected reference alignment using MAFFT (*mafft --add -- keeplength*), based on the user-selected focal supergroup. The extended alignment is then used for phylogenetic placement onto the corresponding reference tree using pplacer likelihood (ML) mode. Placement results are stored as SQLite databases. CSI-SSU uses the script *summarize_results*.*py* to extract and summarize taxonomic classifications from these databases, generating a structured summary output. For visualization, the script *plot_tree*.*py* is used to generate midpoint-rooted trees with placed SSU sequences highlighted, using the ETE3 toolkit.

To identify putative chimeric sequences in large-scale datasets, CSI-SSU implements VSEARCH v2.30.5 using the command *vsearch --uchime_ref*. As a reference database, we used the PR2 SSU dataset (version 5.1.0) downloaded on September 9th, 2025, and clustered with CD-HIT at 95% similarity (“-c 0.95”). Based on this reference, CSI-SSU identifies putative chimeric sequences among the retrieved SSU sequences. The *plot_tree*.*py* script also incorporates VSEARCH results, highlighting the branches of detected chimeric sequences in the final tree visualization.

In addition to the full mode, CSI-SSU provides a placement mode for pre-extracted SSU sequences and a retrieval mode to extract SSU sequences without phylogenetic placement. In the placement mode, the user supplies a FASTA file containing SSU sequences, and the pipeline proceeds directly to the phylogenetic placement and chimeric sequence identification steps, including MAFFT (*--add*) alignment, pplacer placement, VSEARCH, and result summarization with the scripts *summarize_results*.*py* and *plot_tree*.*py*, as described above. In the retrieval mode, the user supplies a nucleotide assembly. SSU sequences are then identified and collected as detailed above, but phylogenetic placement is skipped. This allows users to take the identified SSU sequences and perform their own analyses.

### Phylogenetic placement-based contamination screening of the P10K database

We implemented the CSI-SSU full mode to screen the complete Protist 10,000 Genomes (P10K) database available as of September 9, 2025 (**Supplementary Information, Table S1**). To do so, we downloaded all assemblies available in P10K for each of the major eukaryotic supergroups represented in the database (i.e., Amoebozoa, Archaeplastida, Excavata, Haptista, Obazoa, and TSAR), totaling 2,960 genome-level assemblies. For each assembly, we performed contamination screening using CSI-SSU full mode, following the steps described above and the guidelines provided in the CSI-SSU GitHub repository (https://github.com/AlexTiceLab/CSI-SSU). Analyses were conducted separately for each supergroup, using the corresponding focal supergroup reference alignment and phylogenetic tree provided with the CSI-SSU tool. The P10K database includes assemblies representing the Cryptista supergroup within the Archaeplastida dataset. Therefore, for Archaeplastida screening, we used CSI-SSU with Archaeplastida as the focal group and accounted for this when parsing and presenting the results.

### SSU and COI Phylogenetic reconstruction

To further investigate Amoebozoa as a case study and validate CSI-SSU, we performed phylogenetic reconstructions of 201 amoebozoan taxa from the P10K database (Supplementary Information, Table S1). This approach allows us to refine taxonomic identifications and to assess the performance of CSI-SSU. We constructed a small subunit ribosomal (18S) dataset considering all 201 amoebozoan assemblies. Further, we constructed a cytochrome c oxidase subunit I (COI) dataset focusing only on the testate amoebae order Arcellinida (Tubulinea), since this marker is well sampled for this lineage as it has been traditionally used for phylogenetics in arcellinids [22, 23]. To construct datasets for the 18S and COI markers, we extracted sequences from these assemblies using similarity searches implemented in BLAST+ v2.16.0+ and a custom Python script (**Supplementary Information - File S1**). For each marker, the script automated the creation of BLAST databases using makeblastdb and performed local blastn searches for each query sequence in a multi-FASTA file against each Amoebozoa genomic data. Specifically, the script executed the commands *makeblastdb -in P10KID*.*fasta -dbtype nucl* and *blastn -query marker_query -db P10K*.*fasta -outfmt [script_default_choice]* (**Supplementary Information - File S1**). Aiming for a manageable number of SSU sequences for manual and visual curation, the script retrieved the top five hits per genomic data file, extracting each aligned region along with 1,000 bp of upstream and downstream flanking sequence. This approach enabled recovery of extended SSU and COI regions suitable for downstream phylogenetic analyses and compatible with Amoebozoa SSU and COI data available in the PR^2^ database [24] and NCBI. The orientation of each retrieved sequence was assessed, and sequences were reverse complemented when necessary to match the strand of the original query. For the query sequences, we used 18S data from the PR^2^ database for three representative species of the major amoebozoan clades: *Arcella vulgaris* WP (Tubulinea; GenBank: HM853762.1), *Dictyostelium discoideum* (Evosea; GenBank: AM168040.1), and *Acanthamoeba castellanii* Neff (Discosea; GenBank: U07416.1). As the COI query sequence, we considered the sequence of *Arcella uspiensis* (SRR5396453).

To build a phylogenetically informative datasets, we combined the retrieved sequences with previously published 18S and COI datasets for Amoebozoa, as well as sequences from the PR^2^ database for SSU [13, 15, 23, 24]. For SSU, we curated a non-redundant dataset broadly representative of the major lineages in Amoebozoa (Tubulinea, Evosea, and Discosea) and for COI a dataset broadly representative of the major lineages in Arcellinida, for both markers excluding environmental sequences. This strategy ensured a robust and interpretable dataset for our phylogenetic framework. From a preliminary phylogenetic reconstruction, we curated the SSU and COI datasets used to generate the main trees in this study. This initial analysis allowed us to identify and remove identical or highly similar sequences that resulted from retrieving the top five BLAST+ hits per genome. It also enabled visual inspection of the tree to detect SSU sequences that either failed to cluster within Amoebozoa or formed unusually long branches. Many of these were short sequences (<200 bp) and were excluded from downstream analyses, while some long branches corresponded to full-length sequences that likely represented contaminants. These were retained for further contamination screening.

For contamination screening, we focused on the 18S dataset. After the initial phylogenetic reconstruction (see *Phylogenetic reconstructions* section), any SSU sequences from P10K genome assemblies that represented long branches or did not branch within the Amoebozoa clade were selected for further analysis. These sequences were subjected to additional BLAST+ searches against the PR^2^ database, a curated SSU resource representing eukaryotic diversity, using our custom script and the same parameters described above (**Supplementary Information - File S1**). To perform this search locally, we downloaded the complete PR^2^ database and used it as the reference for the BLAST+ similarity search. This approach allowed us to retrieve SSU sequences from the PR^2^ database that were similar to those of the putative contaminant eukaryotes and to assign their taxonomic affiliations through subsequent phylogenetic analyses.

All phylogenetic reconstructions were based on multiple sequence alignments (MSAs) generated using MAFFT v7.490 with the E-INS-I algorithm and 1000 refinement iterations. Alignments were produced with the following command: *mafft --genadpair --maxiterate 1000 input*.*fasta > output_aligned*.*fasta*. Automated alignment trimming was performed with trimAl v1.2, using the command *trimal -in input_aligned*.*fasta -out output_aligned_trimmed*.*fasta -keepheader -gt [threshold]*, where the gap threshold was set to 0.3 for 18S and 0.5 for COI. Phylogenetic trees were inferred from the trimmed alignments using the maximum likelihood method implemented in IQ-TREE v2.3.6, with ModelFinder for model selection and node support assessed via 1,000 ultrafast bootstrap replicates and 1,000 SH-aLRT tests. Support values of SH-aLRT/UFBoot ≥ 80/95 are considered indicative of strong support, following the recommendations of the original papers describing the development of these node support measures [31, 32]. The analysis was executed with the command *iqtree2 -s aligned_trimmed*.*fasta -alrt 1000 -bb 1000 -m TEST*.

## Results and discussion

### The CSI-SSU tool

The full mode of CSI-SSU begins with a user-provided nucleotide genomic-level assembly in FASTA format for a target organism (**Figure S1**). Based on the organism’s supergroup, users can select the appropriate reference group from those available in CSI-SSU (Amoebozoa, Excavata, TSAR, Archaeplastida, Cryptista, Haptista, CRuMs, Provora, Obazoa), reflecting the current diversity sampling and taxonomy of the PR2 database. In full mode, the tool performs SSU sequence identification and retrieval, chimeric sequence detection, and phylogenetic placement onto a reference SSU tree for the selected supergroup using pplacer. In addition, CSI-SSU conducts bacterial contamination screening using BUSCO searches. The CSI-SSU tool generates a structured output directory containing intermediate and final files, including a summary folder with comprehensive reports and visualizations. All retrieved SSU sequences are provided for downstream analyses, along with taxonomic assignments and placement confidence values based on the likelihood weight ratio (LWR) from pplacer. The tool also produces an annotated phylogenetic tree in which placed sequences are highlighted using a color scheme indicating both taxonomic assignment and chimeric status. Each sequence is annotated across multiple taxonomic levels (supergroup, order, family, and genus), together with corresponding LWR values, facilitating interpretation and manual inspection. Although no universal LWR threshold is defined, simulation studies have shown that higher LWR values are associated with lower placement error. In our empirical analysis of the P10K dataset using CSI-SSU (see results below), most placements fell within LWR values of 0.9–1, corresponding to high-confidence assignments. Lower LWR values were often associated with shorter sequences (e.g., ∼500 bp) or chimeric sequences, whereas higher-confidence placements typically involved longer sequences (∼1,000 bp) positioned within well-supported clades (e.g., bootstrap support >60) and exhibiting consistently high LWR values across taxonomic levels. It is important to note that the resolution of the reference trees is constrained by the current SSU sampling and by the phylogenetic signal of this single marker. Therefore, interpretation of placement results must take into account the taxonomic representation of the target group (or the group to which the SSU sequence is assigned) within the reference dataset. In addition, assembly complexity, contamination, uneven taxonomic sampling, and intrinsic limitations of SSU-based phylogenetic resolution may vary across lineages and influence placement precision. Accordingly, careful visual inspection of the resulting phylogenetic trees and summary outputs is strongly recommended and is explicitly supported by the tool’s automated organization of results, designed to facilitate this process.

Finally, all identified SSU sequences are compiled into a summary FASTA file (parsed_sequences.fasta), which can be used for downstream analyses, including phylogenetic reconstruction.

In addition to the full workflow, CSI-SSU provides retrieval and placement modes. The retrieval mode performs SSU sequence identification and extraction, along with bacterial BUSCO searches and chimeric sequence detection, but does not include phylogenetic placement. This allows users to recover SSU sequences from assemblies for downstream analyses without placing them. The placement mode performs phylogenetic placement of user-provided SSU sequences. In this mode, users can supply SSU sequences in FASTA format from any source, including primer-based 18S amplification of target organisms or metabarcoding projects. Additionally, after running the full workflow, retrieved SSU sequences identified as contaminants can be reanalyzed using the CSI-SSU placement mode with a reference tree corresponding to the identified contaminant supergroup, enabling more precise taxonomic placement and confirmation.

Benchmarking across eukaryotic supergroups, using 2,960 genomic-level assemblies from P10K (see results below), demonstrated that the tool performs efficiently, with generally low runtime, CPU usage, and memory consumption (**Fig. S2**). Median runtimes ranged from 3.9 to 9.2 minutes, median CPU times from 2.9 to 7.4 minutes, and median memory usage from 332 to 459 MB. While most analyses completed quickly, a subset of datasets required substantially more resources, with runtimes reaching up to 154 minutes and memory usage up to 19.9 GB (**Fig. S2**). These higher resource demands were primarily associated with large genome assemblies, with BUSCO accounting for a significant portion of the computational cost due to its mapping and gene annotation steps. All CSI-SSU analyses were conducted on the Quanah partition of the Texas Tech University High Performance Computing Center (HPCC) RedRaider cluster, using a single thread on one node. The Quanah partition is built on Intel Xeon E5-2695 v4 processors and runs Rocky Linux 9.2/8.10, with 36 cores and 192 GB of memory per node. To test the CSI-SSU tool and demonstrate its compatibility with both Linux and macOS, we also performed analyses on a separate workstation: a Mac mini with an Apple M4 Pro chip, 14 cores (10 performance, 4 efficiency), and 48 GB of RAM.

### Contamination Screening of the P10K Database Using CSI-SSU

CSI-SSU retrieved and phylogenetically placed 13,513 SSU sequences from 2,960 genome-level assemblies representing six eukaryotic supergroups obtained from the P10K database (**Figure 1A–B**). In most assemblies, CSI-SSU recovered multiple SSU sequences and placed with high confidence (pplacer likelihood weight ratio ≥ 0.9) within the target supergroup or non-target supergroups (**Figure 1C**). Across supergroups, Amoebozoa exhibited the highest levels of contamination, whereas Obazoa showed the lowest. This pattern is consistent with differences in data origin: Amoebozoa are largely represented by non-model organisms, many of which are derived from environmental single-cell isolations, while Obazoa include several well-established model systems with genome data typically generated from cultured material and listed as Reference level on NCBI (**Supplementary Information - Table S1**). Based on VSEARCH searches, CSI-SSU identified numerous chimeric SSU sequences across genomic assemblies from all surveyed supergroups (**Figure 1D**). Many contaminant SSU sequences were assigned to a wide range of supergroups (**Figure 1E–I**). Among these, TSAR was the most frequent source of contamination, followed by Obazoa, Archaeplastida, and Amoebozoa. One Archaeplastida assembly contained an SSU sequence assigned to Eukaryota_X and placed within Ancyromonas, a group classified as Eukaryota_X in the PR2 database. A small subset of SSU sequences identified and retrieved was placed with a low confidence (likelihood weight ratio as low as 0.2) and often forming long branches (**Supplemental material – Data S1**). These patterns were associated with short or chimeric sequences, which negatively influences the placement accuracy. Bacterial BUSCO scores further indicate that, although many assemblies contain fewer than 30 of the 116 searched bacterial markers (bacteria_odb12), some assemblies show scores exceeding 60 bacterial BUSCO genes, with a few approaching near-complete bacterial gene representation (**Figure 1J**). This pattern is particularly evident in datasets from Amoebozoa, Archaeplastida, Haptista, and TSAR, suggesting substantial bacterial contamination despite the bacterial decontamination effort implemented by P10K project. It is important to note that the presence of bacterial BUSCO genes does not necessarily indicate bacterial contamination, as the possibility of genuine horizontal gene transfer (HGT) events cannot be excluded. Therefore, bacterial BUSCO scores should be considered a proxy and interpreted accordingly. Overall, the results derived from CSI-SSU highlight both the scale and heterogeneity of contamination across protist genomic assemblies available in P10K, as well as the effectiveness of CSI-SSU as a large-scale screening tool. CSI-SSU provides a robust and efficient first-pass framework for SSU recovery and contamination detection. A detailed summary of CSI-SSU results is presented on **Supplementary Information - Table S1** for each of the 2,960.

**Figure 1.**
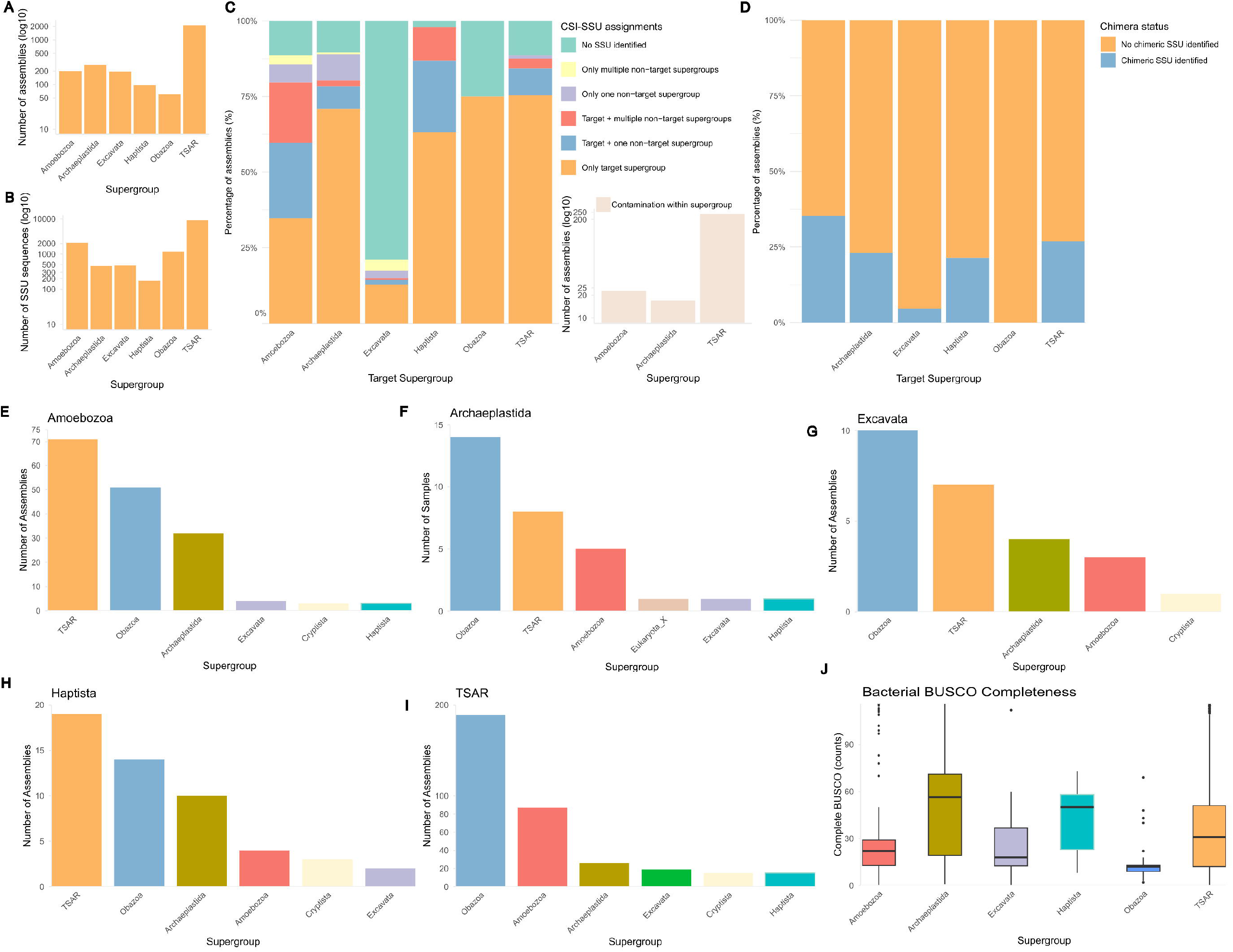
Summary of contamination screening of the P10K database using CSI-SSU tool. **A.** Bar chart showing the total number of assemblies per supergroup in the P10K database screened with CSI-SSU. Values are displayed on a log_10_ scale for improved visualization. **B**. Bar chart showing the total number of SSU sequences identified, retrieved, and placed by CSI-SSU tool per supergroup.**C**. Stacked bar chart showing the percentage of assemblies per supergroup by SSU supergroup assignment category: no SSU identified; SSU sequences assigned only to the target supergroup; SSU sequences assigned to the target and one or multiple non-target supergroups; and SSU sequences assigned only to non-target supergroups (single or multiple). The bar chart shows the total number of assemblies containing within-supergroup contamination. Colors correspond to assignment categories shown in the legend. **D**. Stacked bar chart showing the percentage of assemblies per supergroup with SSU sequences classified as chimeric or non-chimeric. **E**. Bar chart showing the number of Amoebozoa assemblies with SSU sequences assigned to contaminant supergroups. **F**. Bar chart showing the number of Archaeplastida assemblies with SSU sequences assigned to contaminant supergroups. **G**. Bar chart showing the number of Excavata assemblies with SSU sequences assigned to contaminant supergroups. **H**. Bar chart showing the number of Haptista assemblies with SSU sequences assigned to contaminant supergroups. **I**. Bar chart showing the number of TSAR assemblies with SSU sequences assigned to contaminant supergroups. **J**. Boxplot showing the distribution of bacterial markers identified in assemblies from each supergroup. The number of complete BUSCOs represents the count of complete bacterial BUSCO genes detected out of the 116 markers in the bacteria_odb12 database. All displayed results represent assignments with an LWR of at least 0.9 and SSU sequences retrieved and placed with a minimum length of 500 bp.

### Phylogeny-based taxonomic identification of P10K amoebozoan data

Amoebozoan SSU sequences were successfully retrieved from 151 of the 201 genomic datasets available for Amoebozoa in the P10K database (**Supplementary Information - Table S1**). Given the established use of the Cytochrome c oxidase subunit I (COI) marker in Arcellinida phylogenetics, one of the main amoebozoan groups represented in P10K database, we also specifically targeted this marker for arcellinid taxa. Arcellinid COI sequences were successfully recovered from 40 of the 59 genomic data available for Arcellinida (**Supplementary Information - Table S1**). Phylogenetic analyses based on SSU and COI enabled precise taxonomic identification, corroborating CSI-SSU results and supporting most of the original taxonomic assignments in the P10K database (**Supplementary Information - Table S1; Figs. 2 and 3**). They also allowed more accurate classification for 43 taxa, including refined genus- and family-level assignments for taxa previously classified only at higher taxonomic levels. A more precise taxonomic assignment was achieved using more complete SSU and COI datasets curated for Amoebozoa, incorporating not only PR2 sequences but also SSU and COI sequences previously retrieved from transcriptomic data published in the literature but not available on NCBI (**Figs. 2 and 3; Supplementary Information - Table S1**). As species-level identification typically requires extensive morphological and morphometric data in addition to molecular evidence, we adopted a conservative approach and assigned identifications at the genus level based on our phylogenetic results. However, it is worth noting that several genomic data from the P10K database originate from well-established cultures of widely used and shared strains, some originally available in the NCBI database, for which detailed morphological data are available in the literature, allowing confident species-level identifications (**Figs. 2 and 3**; **Supplementary Information - Table S1**).

**Figure 2.**
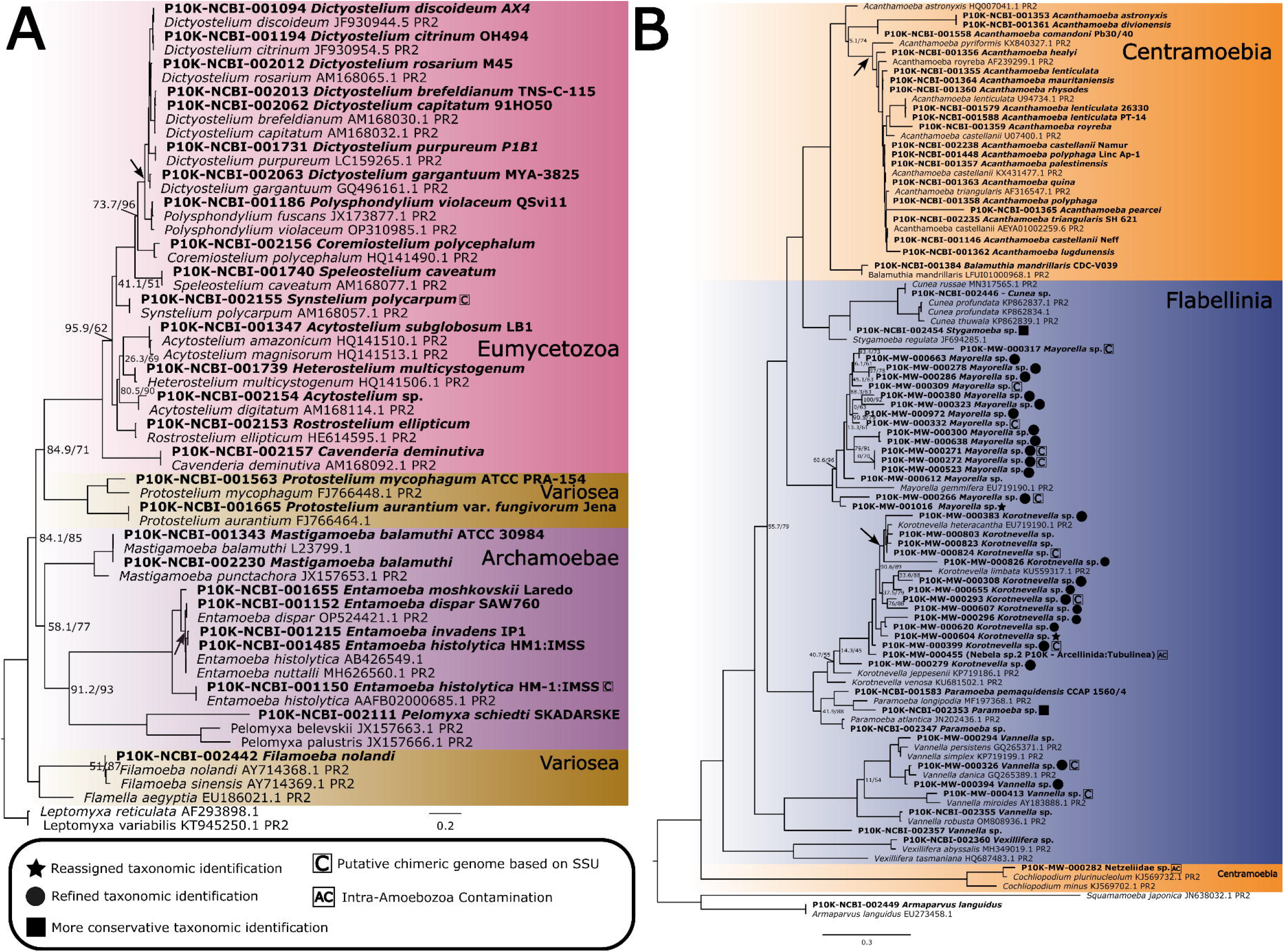
The phylogenetic placement of Amoebozoa genomic data from the P10K focused on the major groups Evosea and Discosea. Maximum-likelihood phylogenetic trees constructed from the Small Subunit ribosomal RNA (SSU) inferred from a subset of the curated dataset presented in Figure S2. **A**. Focuses on the major Amoebozoa group Evosea. Phylogenetic reconstruction was conducted using IQ-TREE v2.3.6, with ModelFinder identifying the best-fit substitution model (TIM2+F+R4). **B**. Focuses on the major Amoebozoa group Discosea. Phylogenetic reconstruction was conducted using IQ-TREE v2.3.6, with ModelFinder identifying the best-fit substitution model (GTR+F+G4). Node support was assessed using both ultrafast bootstrap (UFBoot) and the Shimodaira–Hasegawa approximate likelihood ratio test (SH-aLRT). Support values are reported as SH-aLRT / UFBoot, with values ≥80/95 considered indicative of strong support [32, 33]. For clarity, high-support values are omitted in this figure, as well as support values for nodes above the nodes indicated by the arrows, which are represented mostly by flat branches. The complete tree, including all support values, is shown in Figures S4 and S5. Stars indicate genomic data reassigned to a different taxonomic identity than reported in the P10K database. Filled circles mark indicate cases with refined taxonomic resolution. Filled squares denote more conservative identifications (e.g., genus or family level) rather than the more specific genus or species-level identification originally provided in the P10K database. ‘C’ indicates putative chimeric genomes containing sequences from multiple eukaryotes and ‘AC’ denotes intra-Amoebozoa contamination.

**Figure 3.**
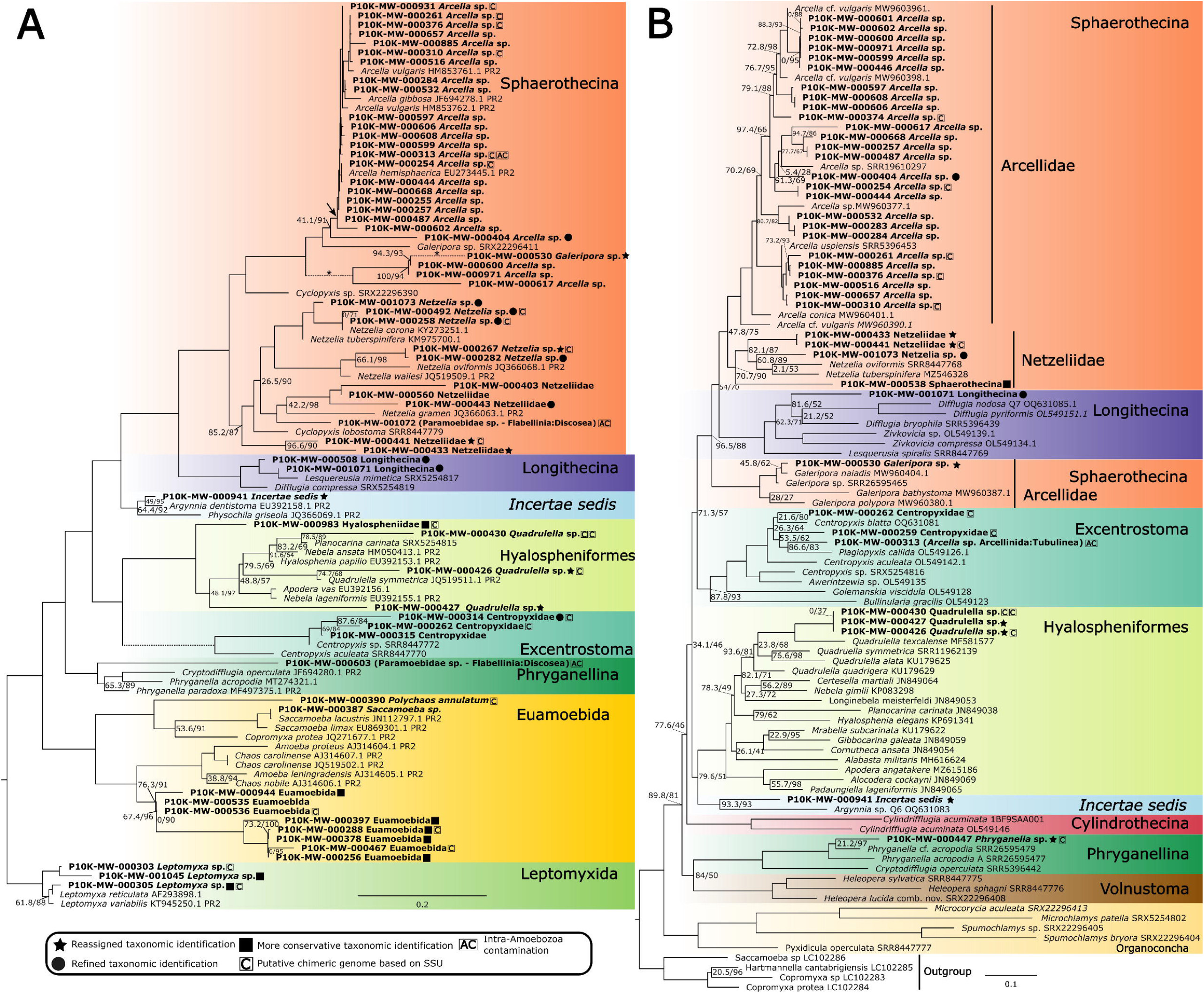
The phylogenetic placement of Amoebozoa genomic data from the P10K focused on the major group Tubulinea. **A.** The maximum-likelihood phylogenetic tree constructed from the Small Subunit ribosomal RNA (SSU) inferred from a subset of the curated dataset presented in Figure S2, focuses on the major Amoebozoa group Tubulinea. Phylogenetic reconstruction was conducted using IQ-TREE v2.3.6, with ModelFinder identifying the best-fit substitution model (TIM3e+G4). The lengths of branches depicted as dashed lines and marked with an asterisk have been reduced by 50% for presentation purposes. **B**. The maximum-likelihood phylogenetic tree of cytochrome c oxidase subunit I (COI) inferred from a curated dataset generated in the present study, focusing on Arcellinida order (Tubulinea:Amoebozoa) comprising COI sequences retrieved from genomes and transcriptomes available in the P10K database, along with reference sequences made available by previous. Phylogenetic reconstruction was conducted using IQ-TREE v2.3.6, with ModelFinder identifying the best-fit substitution model (GTR+F+I+G4). Node support was assessed using both ultrafast bootstrap (UFBoot) and the Shimodaira–Hasegawa approximate likelihood ratio test (SH-aLRT). Support values are reported as SH-aLRT / UFBoot, with values ≥80/95 considered indicative of strong support [32, 33]. For clarity, high-support values are omitted in this figure, as well as support values for nodes above the one indicated by the arrow, which are represented mostly by flat branches. The complete tree, including all support values, is shown in Figures S6 and S7. Stars indicate genomic data reassigned to a different taxonomic identity than reported in the P10K database. Filled circles mark indicate cases with refined taxonomic resolution. Filled squares denote more conservative identifications (e.g., genus or family level) rather than the more specific genus or species-level identification originally provided in the P10K database. ‘C’ indicates putative chimeric genomes containing sequences from multiple eukaryotes and ‘AC’ denotes intra-Amoebozoa contamination.

Notably, seven original taxonomic assignments from the P10K database were not corroborated by our phylogenetic reconstructions (**Figures 2 and 3**; **Supplementary Information - Table S1**). This was most apparent within Arcellinida, where misidentifications primarily involved closely related or morphologically similar genera and families. Although Arcellinida are well known for their test (shell), which provides informative taxonomic characters, convergent evolution has led to similar shell morphologies across distantly related lineages. For example, species of *Difflugia* (Difflugiidae), *Hyalosphenia* (Hyalospheniidae), and *Netzelia* (Netzeliidae) often possess rounded, ovoid, or elongated shells, which can lead to misidentification if other shell features (e.g., aperture shape, composition) or cellular characteristics are not considered. Consistent with this, our analyses revealed that several taxa initially assigned to Difflugiidae (infraorder Longithecina) belong to Hyalospheniidae (Hyalospheniformes) or Netzeliidae (Sphaerothecina) (**Supplementary Information - Table S1**). These infraorders are distantly related, with their last common ancestor estimated to have lived over 500 million years ago [15, 25], making such misassignments evolutionarily significant. By comparison, CSI-SSU correctly placed arcellinid SSU sequences within Arcellinida with high confidence (LWR ≥ 0.9, most LWR = 1), consistent with results from SSU and COI phylogenetic reconstruction (**Supplementary Information - Table S1; DataS1**). Outside Arcellinida, only one notable case of misidentification was observed: *Pessonella* sp. PRA-29, a culture originally submitted to the American Type Culture Collection (ATCC) under the genus *Pessonella*, was later described as a new genus and species, *Armaparvus languidus*, representing the correct taxonomic identification for this organism [26].

### Contamination screening of the P10K amoebozoan data

Non-amoebozoan SSU sequences were retrieved from 58 of the 201 genomic datasets available for Amoebozoa in the P10K database (**Supplementary Information - Table S1**). The phylogenetic analysis of these non-amoebozoan SSU and SSU sequences from the PR^2^ database reveals a widespread contamination of the P10K amoebozoan dataset by a taxonomically diverse set of eukaryotic lineages, mirroring the ecological complexity of the environments these protists inhabit (**Figure 3 and Supplementary Information - Table S1**). Among the contaminant groups were fungi and metazoans, including sequences from arthropods and nematodes, likely introduced via soil particles, organic debris, or sample handling procedures (**Figure 4**). SSU sequences affiliated with ciliates were particularly abundant, with representatives spanning multiple clades such as Vorticellidae, *Coleps*, and *Pseudomicrothorax* (**Figure 4**). This reflects the high diversity and abundance of ciliates across environments, leading to their consistent identification as contaminants.

**Figure 4.**
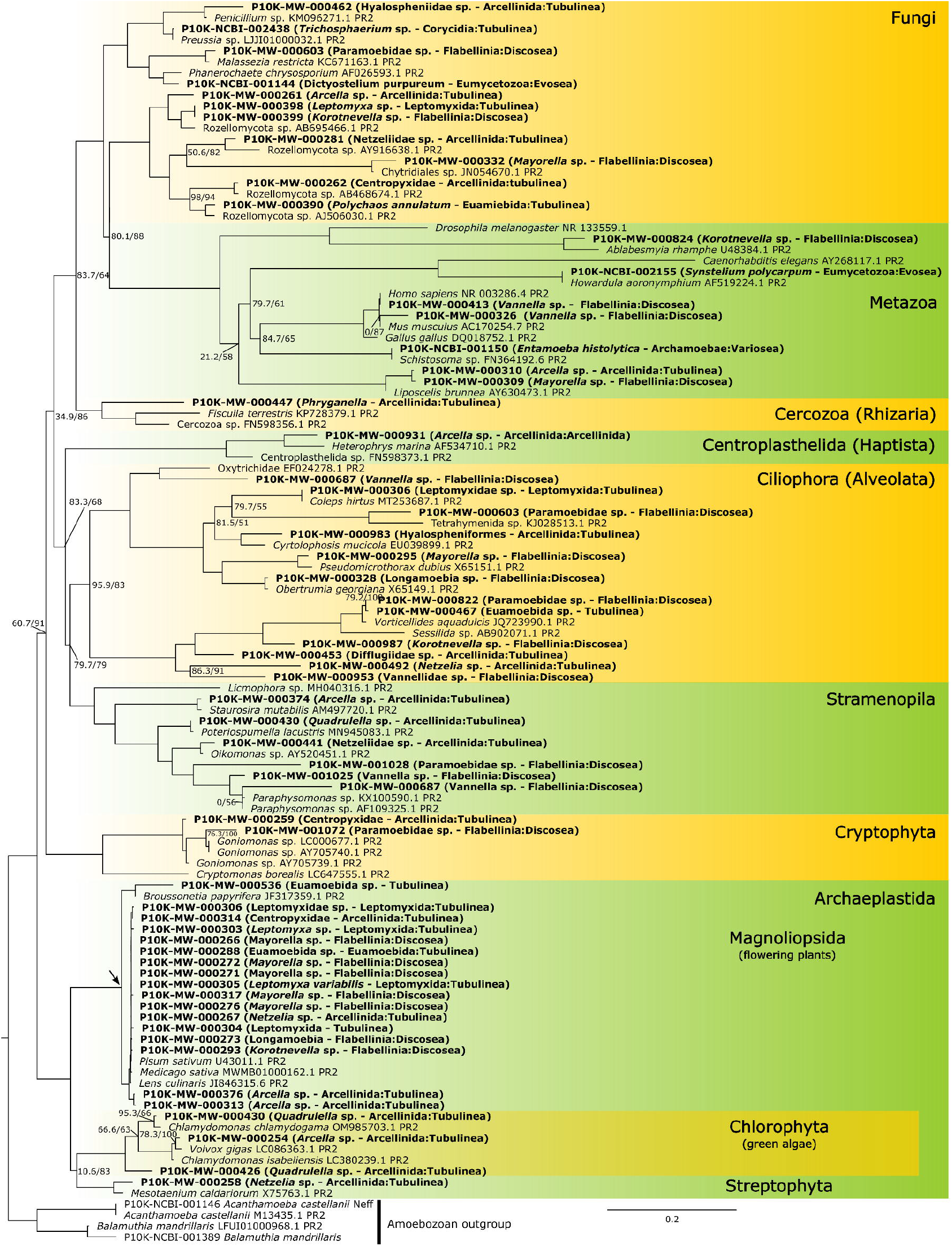
Contamination-screening phylogenetic tree. The maximum-likelihood phylogenetic tree constructed from the Small Subunit ribosomal RNA (SSU), inferred from a curated dataset generated in the present study, comprises SSU sequences of putative non-Amoebozoan contaminants retrieved from genomes and transcriptomes available in the P10K database, along with reference sequences from the PR^2^ database (indicated by PR^2^ IDs), identified through BLAST+ similarity searches using the putative contaminant SSU sequences as queries. Phylogenetic reconstruction was conducted using IQ-TREE v2.3.6, with ModelFinder identifying the best-fit substitution model (TN+F+I+G4). Node support was assessed using both ultrafast bootstrap (UFBoot) and the Shimodaira–Hasegawa approximate likelihood ratio test (SH-aLRT). Support values are reported as SH-aLRT / UFBoot, with values ≥80/95 considered indicative of strong support [32, 33]. For clarity, high-support values are omitted in this figure, as well as support values for nodes above the one indicated by the arrow, which are represented mostly by flat branches. The complete tree, including all support values, is shown in Figure S8. The taxonomic identification of Amoebozoa taxa corresponding to the displayed P10K ID codes is shown in parentheses.

Additional contaminants included lineages within the Stramenopiles such as *Paraphysomonas* and *Poterioochromonas*, as well as centrohelid heliozoans (Haptista) and cercozoans (Rhizaria) (**Figure 4**). Several photosynthetic eukaryotes represented another major group of contaminants, including land plants, especially angiosperms of the Fabaceae family, likely introduced through pollen or plant debris, as well as green algae from the Chlorophyceae (e.g., *Chlamydomonas, Hyalomonas*) and Zygnematophyceae, the closest relatives of land plants (**Figure 4**). It is worth noting that symbiotic associations may also contribute to the high levels of apparent contamination observed in Amoebozoa and other eukaryotic groups [27]. For example, several taxa of testate amoebae are known to be consistently associated with green algae such as Chlorella (Chlorophyta) [27]. The identification of some contaminant sequences may therefore reflect biological associations rather than external contamination. By comparison, CSI-SSU assignments were consistent with the phylogenetic reconstruction results (**Supplementary Information - Table S1**). Notably, for the phylogenetic reconstruction approach we implemented, only the top five BLAST hits were retained. In contrast, CSI-SSU retrieves all sequences identified by BLAST hits, resulting in a more comprehensive identification and classification of contaminant SSU sequences. This is particularly important for assemblies contaminated by multiple non-target supergroups or containing multiple SSU copies per group (**Figure 1; Supplementary Information - Table S1; Data S1**).

### Quality assessment of the P10K data

Collectively, CSI-SSU and the comparative reconstruction approach findings corroborates that contamination in eukaryotic assemblies is both frequent and taxonomically widespread, encompassing multiple branches of the eukaryotic tree. Based on our contamination screening results and other parameter, we can evaluated the current state of microbial eukaryote genomic data in the P10K database. Samples can be considered relatively higher quality if they contain the SSU marker for the target organisms, show no evidence of contamination based on SSU phylogenetic reconstruction, and have a eukaryotic BUSCO completeness score of at least 50% (**Supplementary Information - Table S1**). The eukaryotic BUSCO scores, already included in the P10K metadata, were used here to assess completeness. Such samples are more likely to yield meaningful results in downstream analyses, including phylogenomics and comparative genomics (**Supplementary Information - Table S1**). It is important to note that SSU-based contamination screening has limitations and cannot detect all sources of potential contamination. Specifically, while the presence of a contaminant SSU informs the chimeric characteristics of a given assembly, the absence of SSU sequences does not necessarily indicate that a sample is free of contamination, as other contaminant sequences may still be present. While we are considering a relatively low BUSCO threshold (≥50%) to include a broader range of genomic data, more stringent completeness cutoffs and additional quality metrics, such as genome contiguity, N50, and assembly size, may be required depending on the intended analyses. Similarly, P10K samples lacking key markers, flagged as contaminated by SSU phylogenetic inference, or with BUSCO scores below 50% represent lower-quality or incomplete genomic data (**Supplementary Information - Table S1**). Several assemblies require decontamination or additional sequencing prior to downstream analyses (**Supplementary Information - Table S1**). This assessment also highlights that several major lineages remain unsampled (or poorly sampled) at the genomic level and provides a guide for targeted sequencing efforts aimed at expanding taxonomic coverage in the P10K database.

Contamination in protist genomic data, is expected and widespread due to their ecological context: these organisms inhabit complex microbial communities, engage in predation, and often form close associations with other microbes, making isolation of pure genomic material challenging [13, 28, 29]. Contaminant sequences can compromise downstream analyses by inflating genome complexity, distorting gene family evolution, misleading functional annotation, and generating artifactual phylogenetic relationships [4, 30, 31]. While cultivation and single-cell isolation techniques, combined with careful sterile handling, filtration, and repeated transfers, can reduce contamination, complete elimination is rarely possible, particularly for environmentally derived specimens. Therefore, comprehensive contamination screening, such as single-marker phylogenetic analysis (e.g., SSU) coupled with genome-level examination, is essential for accurate data curation and taxonomic validation. Additionally, the P10K database currently lacks detailed metadata and specimen documentation, including voucher images and information on sequencing platforms, which limits reproducibility, taxonomic verification, and interpretation of genomic data. Incorporating such metadata, alongside rigorous contamination screening, would enhance the reliability, usability, and long-term scientific value of microbial eukaryotic genomic resources.

## Conclusions

This study introduces CSI-SSU, a scalable and reproducible tool for phylogenetically informed contamination screening and taxonomic validation of genomic datasets, and demonstrates its performance across the microbial eukaryotic assemblies from the P10K database. By integrating SSU-based sequence retrieval, chimeric detection, and phylogenetic placement within a unified workflow, CSI-SSU enables rapid identification of target lineages while simultaneously detecting non-target eukaryotic and bacterial contamination. Applied at scale, CSI-SSU tool revealed widespread and taxonomically diverse contamination, reflecting the ecological complexity of protist-associated environments and the inherent difficulty of generating contamination-free genomic data. Importantly, our results highlight that single-marker phylogenetic screening, especially using SSU given its traditional marker and existence of curated databases, sampling in diverse, provides a robust and efficient strategy for both taxonomic validation and exploratory contamination detection. CSI-SSU offers a critical entry point for large-scale data curation, allowing researchers to rapidly assess dataset quality and prioritize assemblies for downstream analyses. By improving the accuracy and interpretability of genomic resources, CSI-SSU contributes directly to enabling reliable evolutionary, ecological, and functional genomic studies. More broadly, it establishes a generalizable framework for curating the rapidly expanding volume of microbial eukaryotic genomic data. As initiatives like P10K continue to grow, integrating CSI-SSU tool into standard pipelines, alongside improved metadata, contamination control strategies, and reproducible workflows, will be essential to maximize the scientific value of these resources and to support robust inferences about the evolution and diversification of eukaryotic life.

## Availability of data and materials

The CSI-SSU tool is hosted on GitHub (https://github.com/AlexTiceLab/CSI-SSU/). All compiled and curated SSU and COI datasets associated with this manuscript are publicly available on FigShare at https://doi.org/10.6084/m9.figshare.29814947.

## Competing interests

The authors declare that they have no competing interests

## Funding

A.L.P-S and R.E.J were supported by startup funds provided to A.K.T. by Texas Tech University. This work was supported by the National Science Foundation Division of Environmental Biology (2100888) awarded to M.W.B., D.J.G.L. is supported by a FAPESP award #2019/22815-2.

## Authors’ contributions

**Alfredo L. Porfirio-Sousa:** Conceptualization, Data curation, Visualization, Formal analysis, Investigation, Writing – original draft, Writing – review & editing. **Robert E. Jones**: Investigation, Data curation, Writing – review & editing. **Matthew W. Brown:** Investigation, Writing – review & editing. **Daniel Lahr:** Investigation, Writing – review & editing. **Alexander K. Tice:** Conceptualization, Data curation, Visualization, Formal analysis, Investigation, Writing – original draft, Writing – review & editing, Resources, Supervision, Funding acquisition.

## Acknowledgments

The authors acknowledge the High Performance Computing Center (HPCC) at Texas Tech University for providing computational resources that have contributed to the research results reported within this paper (URL: http://www.hpcc.ttu.edu). We are grateful to Dr. Carl Seaquist for testing the CSI-SSU tool.

## Declaration of generative AI in scientific writing

During the preparation of this work the authors used ChatGPT GPT-4o to improve the readability and language of the first draft of the manuscript. After using this tool, the authors reviewed and edited the content as needed and take full responsibility for the content of the published article.

## Ethics approval and consent to participate

Not applicable

## Consent for publication

Not applicable

## Notes

https://doi.org/10.6084/m9.figshare.29814947

### Competing Interest Statement

The authors have declared no competing interest.

### Summary of Updates

The new version of the manuscript presents the CSI-SSU tool, which we developed to identify contaminant sequences based on SSU (18S) data. The tool is fully available on GitHub and provides an effective and efficient approach for contamination screening. Using CSI-SSU, we expanded our analysis to the entire P10K database, covering all represented eukaryotic supergroups. In total, we screened 2,960 genomes and identified putative contaminants based on SSU sequences, including chimeric SSUs. For Amoebozoa, we further provide phylogenetic reconstructions that corroborate the CSI-SSU results and enable more precise taxonomic identification of amoebozoan sequences within the P10K database. Importantly, the key findings and conclusions from the original submission are supported by the CSI-SSU results and are now comprehensively validated across all six eukaryotic supergroups represented in the P10K database.

## References

1. Lewin HA, Robinson GE, Kress WJ, Baker WJ, Coddington J, Crandall KA, et al. Earth BioGenome Project: Sequencing life for the future of life. Proceedings of the National Academy of Sciences. 2018;115:4325–33. 10.1073/pnas.1720115115.

2. Gao X, Chen K, Xiong J, Zou D, Yang F, Ma Y, et al. The P10K database: a data portal for the protist 10 000 genomes project. Nucleic Acids Research. 2024;52:D747–55. 10.1093/nar/gkad992.

3. Brown MW, Tice AK. A genetic toolbox for marine protists. Nat Methods. 2020;17:469–70. 10.1038/s41592-020-0794-z.

4. Schoenle A, Francis O, Archibald JM, Burki F, Vries J de, Dumack K, et al. Protist genomics: key to understanding eukaryotic evolution. Trends in Genetics. 2025;0. 10.1016/j.tig.2025.05.004.

5. Burki F, Roger AJ, Brown MW, Simpson AGB. The New Tree of Eukaryotes. Trends in Ecology & Evolution. 2020;35:43–55. 10.1016/j.tree.2019.08.008.

6. Williamson K, Eme L, Baños H, McCarthy CGP, Susko E, Kamikawa R, et al. A robustly rooted tree of eukaryotes reveals their excavate ancestry. Nature. 2025;640:974–81. 10.1038/s41586-025-08709-5.

7. López-García P, Moreira D. The Syntrophy hypothesis for the origin of eukaryotes revisited. Nat Microbiol. 2020;5:655–67. 10.1038/s41564-020-0710-4.

8. Keeling PJ, Burki F, Wilcox HM, Allam B, Allen EE, Amaral-Zettler LA, et al. The Marine Microbial Eukaryote Transcriptome Sequencing Project (MMETSP): Illuminating the Functional Diversity of Eukaryotic Life in the Oceans through Transcriptome Sequencing. PLOS Biology. 2014;12:e1001889. 10.1371/journal.pbio.1001889.

9. The Darwin Tree of Life Project Consortium. Sequence locally, think globally: The Darwin Tree of Life Project. Proceedings of the National Academy of Sciences. 2022;119:e2115642118. 10.1073/pnas.2115642118.

10. Leebens-Mack JH, Barker MS, Carpenter EJ, Deyholos MK, Gitzendanner MA, Graham SW, et al. One thousand plant transcriptomes and the phylogenomics of green plants. Nature. 2019;574:679–85. 10.1038/s41586-019-1693-2.

11. Lahr DJ. An emerging paradigm for the origin and evolution of shelled amoebae, integrating advances from molecular phylogenetics, morphology and paleontology. Mem Inst Oswaldo Cruz. 2021;116:e200620. 10.1590/0074-02760200620.

12. Ribeiro GM, Lahr DJG. Survival in a Changing World: The role of transcriptomics and the urgent need for genomes to understand Arcellinida’s adaptive capabilities. Acta Protozoologica. 2025;2024 Volume 63, Special Issue / Early View.

13. Kang S, Tice AK, Spiegel FW, Silberman JD, Pánek T, Cepicka I, et al. Between a Pod and a Hard Test: The Deep Evolution of Amoebae. Molecular Biology and Evolution. 2017;34:2258–70. 10.1093/molbev/msx162.

14. Onsbring H, Tice AK, Barton BT, Brown MW, Ettema TJG. An efficient single-cell transcriptomics workflow for microbial eukaryotes benchmarked on Giardia intestinalis cells. BMC Genomics. 2020;21:448. 10.1186/s12864-020-06858-7.

15. Porfirio-Sousa AL, Tice AK, Morais L, Ribeiro GM, Blandenier Q, Dumack K, et al. Amoebozoan testate amoebae illuminate the diversity of heterotrophs and the complexity of ecosystems throughout geological time. Proceedings of the National Academy of Sciences. 2024;121:e2319628121. 10.1073/pnas.2319628121.

16. Parks DH, Imelfort M, Skennerton CT, Hugenholtz P, Tyson GW. CheckM: assessing the quality of microbial genomes recovered from isolates, single cells, and metagenomes. Genome Res. 2015;25:1043–55. 10.1101/gr.186072.114.

17. Astashyn A, Tvedte ES, Sweeney D, Sapojnikov V, Bouk N, Joukov V, et al. Rapid and sensitive detection of genome contamination at scale with FCS-GX. Genome Biol. 2024;25:60. 10.1186/s13059-024-03198-7.

18. Camacho C, Coulouris G, Avagyan V, Ma N, Papadopoulos J, Bealer K, et al. BLAST+: architecture and applications. BMC Bioinformatics. 2009;10:421. 10.1186/1471-2105-10-421.

19. Katoh K, Toh H. Recent developments in the MAFFT multiple sequence alignment program. Brief Bioinform. 2008;9:286–98. 10.1093/bib/bbn013.

20. Matsen FA, Kodner RB, Armbrust EV. pplacer: linear time maximum-likelihood and Bayesian phylogenetic placement of sequences onto a fixed reference tree. BMC Bioinformatics. 2010;11:538. 10.1186/1471-2105-11-538.

21. PeerJ VSEARCH: a versatile open source tool for metagenomics. https://peerj.com/articles/2584/#MainContent. Accessed 23 Mar 2026.

22. Kosakyan A, Heger TJ, Leander BS, Todorov M, Mitchell EAD, Lara E. COI Barcoding of Nebelid Testate Amoebae (Amoebozoa: Arcellinida): Extensive Cryptic Diversity and Redefinition of the Hyalospheniidae Schultze. Protist. 2012;163:415–34. 10.1016/j.protis.2011.10.003.

23. Ribeiro GM, Useros F, Dumack K, González-Miguéns R, Siemensma F, Porfírio-Sousa AL, et al. Expansion of the cytochrome C oxidase subunit I database and description of four new lobose testate amoebae species (Amoebozoa; Arcellinida). European Journal of Protistology. 2023;91:126013. 10.1016/j.ejop.2023.126013.

24. Guillou L, Bachar D, Audic S, Bass D, Berney C, Bittner L, et al. The Protist Ribosomal Reference database (PR2): a catalog of unicellular eukaryote Small Sub-Unit rRNA sequences with curated taxonomy. Nucleic Acids Research. 2013;41:D597–604. 10.1093/nar/gks1160.

25. Lahr DJG, Kosakyan A, Lara E, Mitchell EAD, Morais L, Porfirio-Sousa AL, et al. Phylogenomics and Morphological Reconstruction of Arcellinida Testate Amoebae Highlight Diversity of Microbial Eukaryotes in the Neoproterozoic. Current Biology. 2019;29:991–1001.e3. 10.1016/j.cub.2019.01.078.

26. Schuler GA, Brown MW. Description of Armaparvus languidus n. gen. n. sp. Confirms Ultrastructural Unity of Cutosea (Amoebozoa, Evosea). Journal of Eukaryotic Microbiology. 2019;66:158–66. 10.1111/jeu.12640.

27. Gomaa F, Kosakyan A, Heger TJ, Corsaro D, Mitchell EAD, Lara E. One Alga to Rule them All: Unrelated Mixotrophic Testate Amoebae (Amoebozoa, Rhizaria and Stramenopiles) Share the Same Symbiont (Trebouxiophyceae). Protist. 2014;165:161–76. 10.1016/j.protis.2014.01.002.

28. Kosakyan A, Gomaa F, Lara E, Lahr DJG. Current and future perspectives on the systematics, taxonomy and nomenclature of testate amoebae. European Journal of Protistology. 2016;55:105–17. 10.1016/j.ejop.2016.02.001.

29. Burki F, Sandin MM, Jamy M. Diversity and ecology of protists revealed by metabarcoding. Current Biology. 2021;31:R1267–80. 10.1016/j.cub.2021.07.066.

30. Francois CM, Durand F, Figuet E, Galtier N. Prevalence and Implications of Contamination in Public Genomic Resources: A Case Study of 43 Reference Arthropod Assemblies. G3 (Bethesda). 2020;10:721–30. 10.1534/g3.119.400758.

31. Cornet L, Baurain D. Contamination detection in genomic data: more is not enough. Genome Biol. 2022;23:60. 10.1186/s13059-022-02619-9.

32. Guindon S, Dufayard J-F, Lefort V, Anisimova M, Hordijk W, Gascuel O. New Algorithms and Methods to Estimate Maximum-Likelihood Phylogenies: Assessing the Performance of PhyML 3.0. Syst Biol. 2010;59:307–21. 10.1093/sysbio/syq010.

33. Minh BQ, Nguyen MAT, von Haeseler A. Ultrafast Approximation for Phylogenetic Bootstrap. Mol Biol Evol. 2013;30:1188–95. 10.1093/molbev/mst024.

