## Supplementary Information for "CSI-SSU: Phylogenetic contamination screening of genomic datasets, demonstrated on the Protist 10,000 Genomes (P10K) database"

**This PDF file includes:**

Table S1 legend

File S1 legend

Supplementary Figures S1 – S8

Supplementary dataset legends (Data S1 – S8)

**Table S1**. Summary of genomic-level data from the Protist 10,000 Genomes (P10K) initiative, including all represented supergroups (Amoebozoa, Archaeplastida, Excavata, Haptista, Obazoa, and TSAR). The table includes species names, sample and assembly identifiers, contamination assessment based on the CSI-SSU tool, bacterial BUSCO results, chimeric SSU identification, and basic assembly and annotation metrics. Assembly quality metrics comprise total assembly size (Mb), number of scaffolds or transcripts, N50 (bp), genome completeness (% BUSCO), predicted gene count, CDS completeness (percentage of genes with both start and stop codons), and final annotation quality level (High, Medium, Low). These metrics were not generated in the present study but are reported as provided in the P10K metadata (Gao et al., 2024). In addition to CSI-SSU results, the table includes, for Amoebozoa, the detection of SSU markers (and COI for Arcellinida) and phylogenetically informed taxonomic identification based on SSU and COI phylogenetic reconstructions.

**File S1.** Custom Python script used to automate local BLAST+ searches across a folder of genome or transcriptome FASTA files. For each input query sequence, the script retrieves the top *N* high-scoring hits per target file. Optionally, it can extract user-defined flanking regions around each hit, reorient the retrieved sequences to match the query strand, and save the output in FASTA format. This tool facilitates high-throughput retrieval of target regions from large-scale genomic datasets through similarity search and was used in this study to extract targeted SSU and COI sequences from assembled genomes and transcriptomes of Amoebozoa available in the P10K database.

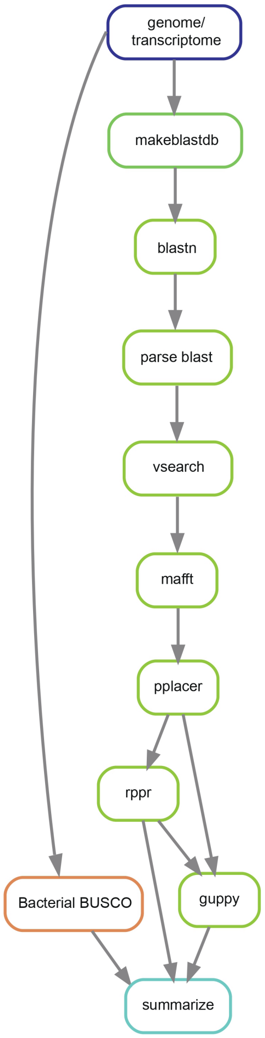


**Figure S1. Overview of the CSI-SSU tool workflow.** CSI-SSU is a command-line pipeline for screening SSU (18S rRNA) sequences from genomic and transcriptomic assemblies. BLAST+ v2.17.0 databases are generated from input assemblies (makeblastdb) and queried using reference SSU sequences (blastn). Hits are parsed to retrieve and orient candidate SSU sequences, which are then screened for chimeric sequences using VSEARCH v2.30.5, aligned to one of nine curated reference alignments with MAFFT v7.526, and phylogenetically placed onto reference trees using pplacer v1.1.alpha20. These reference alignments and trees were constructed from the 18S dataset of the Protist Ribosomal Reference (PR2) database and represent the nine major eukaryotic supergroups (Amoebozoa, Excavata, TSAR, Archaeplastida, Cryptista, Haptista, CRuMs, Provora, and Obazoa). Placement results are processed with guppy and rppr and summarized to generate taxonomic assignments. In parallel, bacterial contamination is assessed using BUSCO. Final outputs integrate phylogenetic placement, contamination screening, and summary statistics

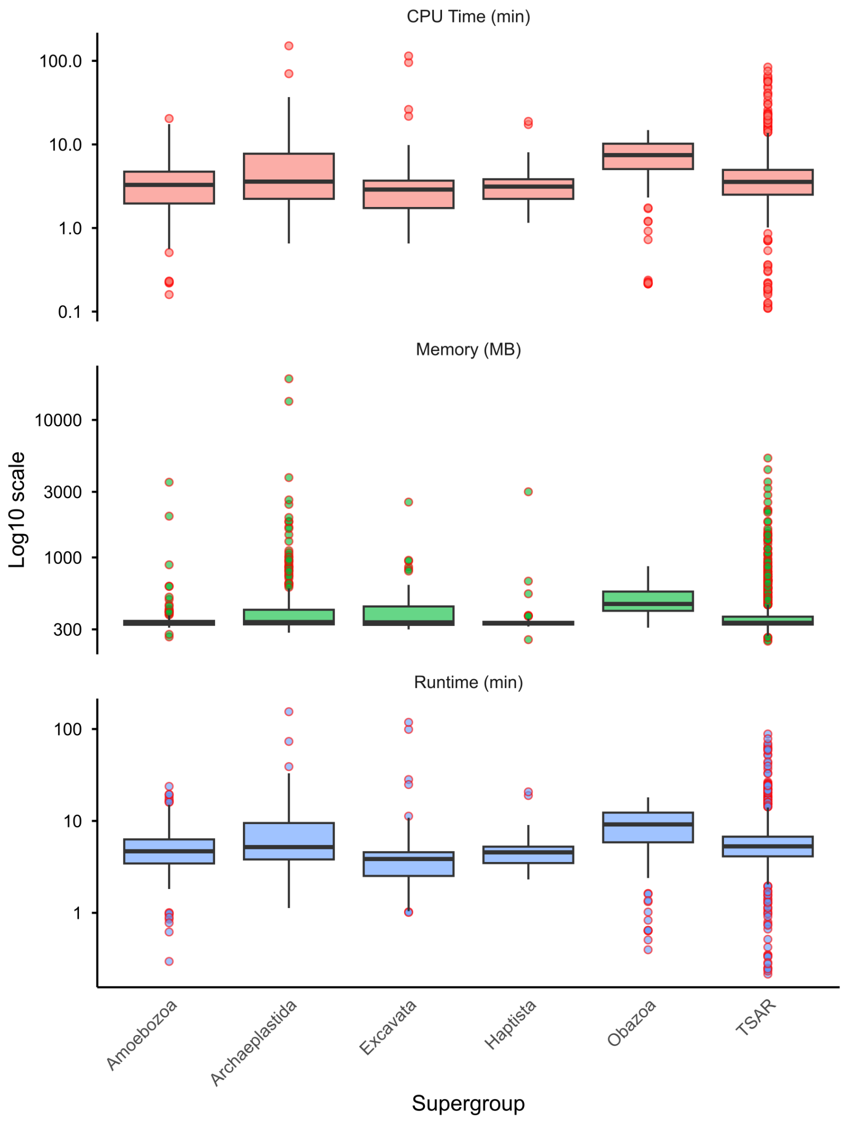


**Fig. S2. Benchmarking performance of CSI-SSU across major eukaryotic supergroups.** Boxplots summarize CPU time in minutes (top), memory usage in megabyte (middle), and total runtime in minutes (bottom) for analyses of 2,960 genomic-level assemblies from the P10K dataset. Values are shown on a $\log_{10}$ scale. Boxes represent the interquartile range (IQR) with median values indicated by horizontal lines; whiskers extend to 1.5× IQR. Individual points denote outliers. Across supergroups (Amoebozoa, Archaeplastida, Excavata, Haptista, Obazoa, and TSAR), most runs show low and consistent computational requirements, while a subset of analyses exhibit higher resource usage. The analyses were performed on the Quanah partition of the Texas Tech University High Performance Computing Center (HPCC) RedRaider cluster. The Quanah partition is built on Intel Xeon E5-2695 v4 processor and runs Rocky Linux 9.2/8.10, with 36 cores and 192 GB of memory available per node. All analyses were conducted using a single thread on one node of the Quanah partition.

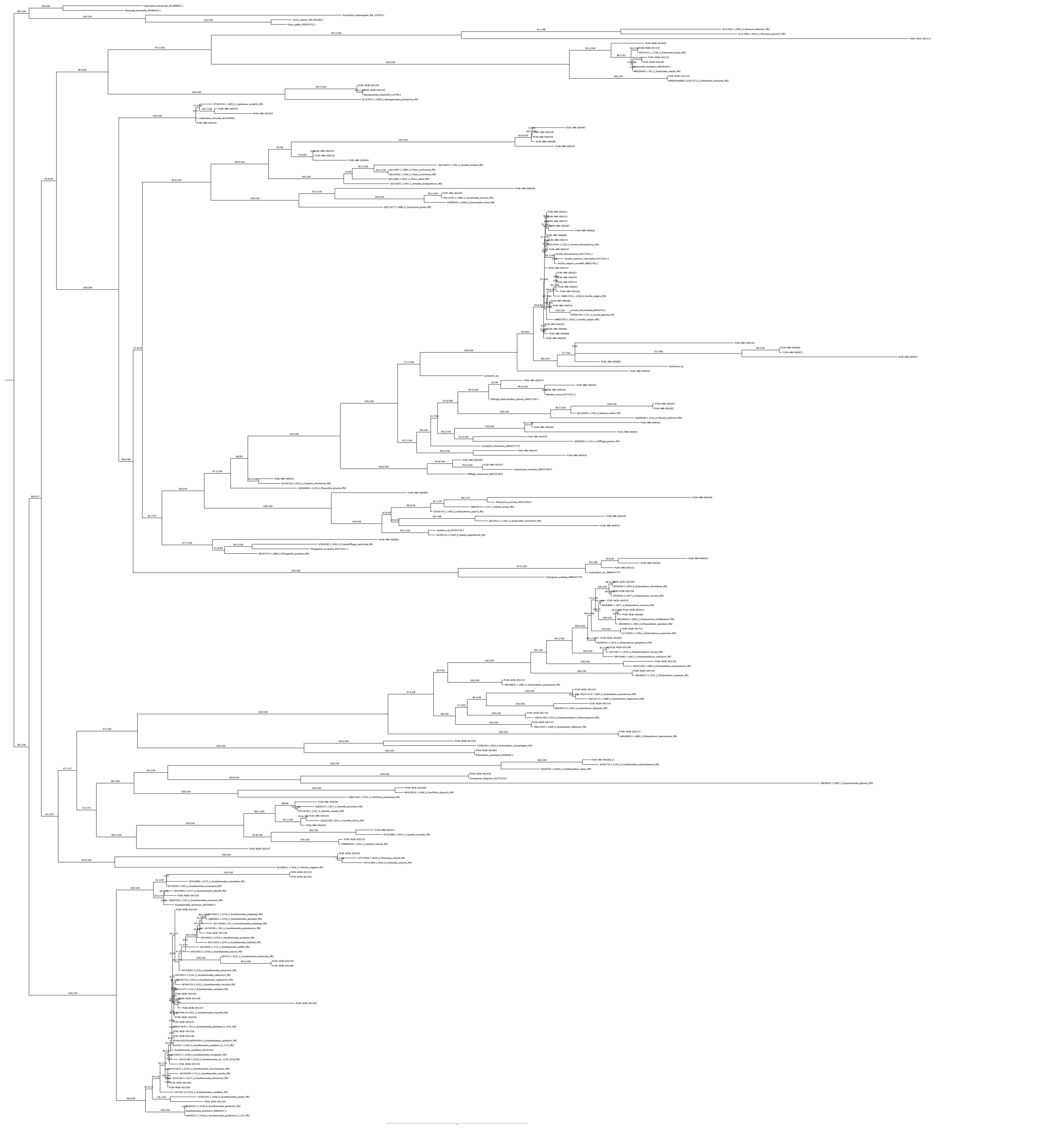

**Figure S3.** The maximum-likelihood phylogenetic tree of Small Subunit ribosomal RNA (SSU) inferred from a curated dataset generated in the present study, comprising SSU sequences retrieved from genomes and transcriptomes available in the P10K database, along with reference sequences from the PR$^2$ database (indicated by PR$^2$ IDs) and NCBI. The dataset includes representatives from the three major Amoebozoa groups: Tubulinea, Evosea, and Discosea. Subsets of this dataset were used to generate the trees presented in Figures 2 and 3A, and Figures S4–S7. Phylogenetic reconstruction was conducted using IQ-TREE v2.3.6, with ModelFinder identifying the best-fit substitution model (GTR+F+I+G4). Node support was assessed using both ultrafast bootstrap (UFBoot) and the Shimodaira–Hasegawa approximate likelihood ratio test (SH-aLRT). Support values are reported as SH-aLRT / UFBoot, with values ≥80/95 considered indicative of strong support.

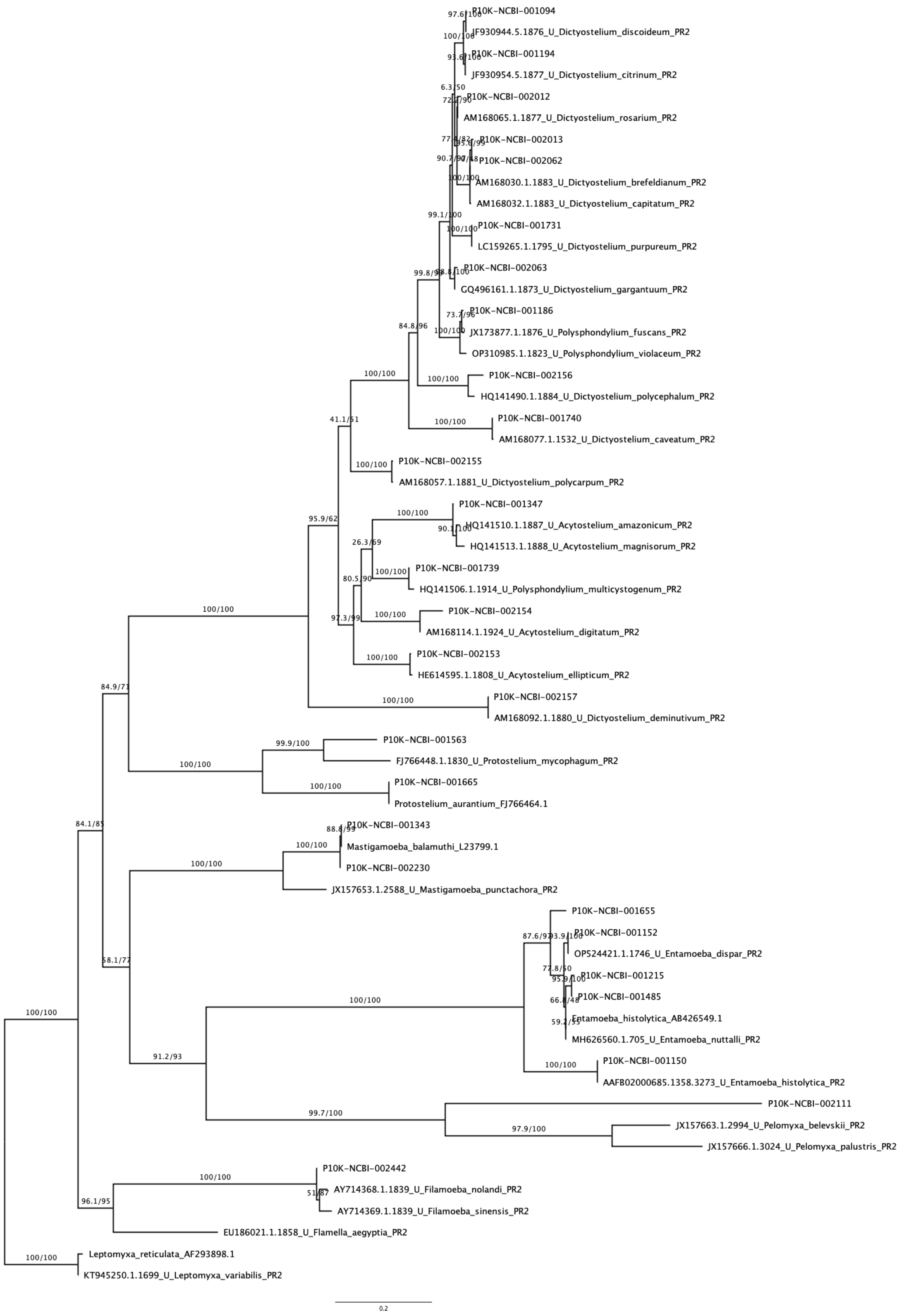


**Figure S4.** The maximum-likelihood phylogenetic tree of Small Subunit ribosomal RNA (SSU) inferred from a subset of the curated dataset presented in Figure S3, focuses on the major Amoebozoa group

Evosea and is shown as Figure 2A in the main text of the manuscript. Phylogenetic reconstruction was conducted using IQ-TREE v2.3.6, with ModelFinder identifying the best-fit substitution model (TIM2+F+R4). Node support was assessed using both ultrafast bootstrap (UFBoot) and the Shimodaira–Hasegawa approximate likelihood ratio test (SH-aLRT). Support values are reported as SH-aLRT / UFBoot, with values ≥80/95 considered indicative of strong support.

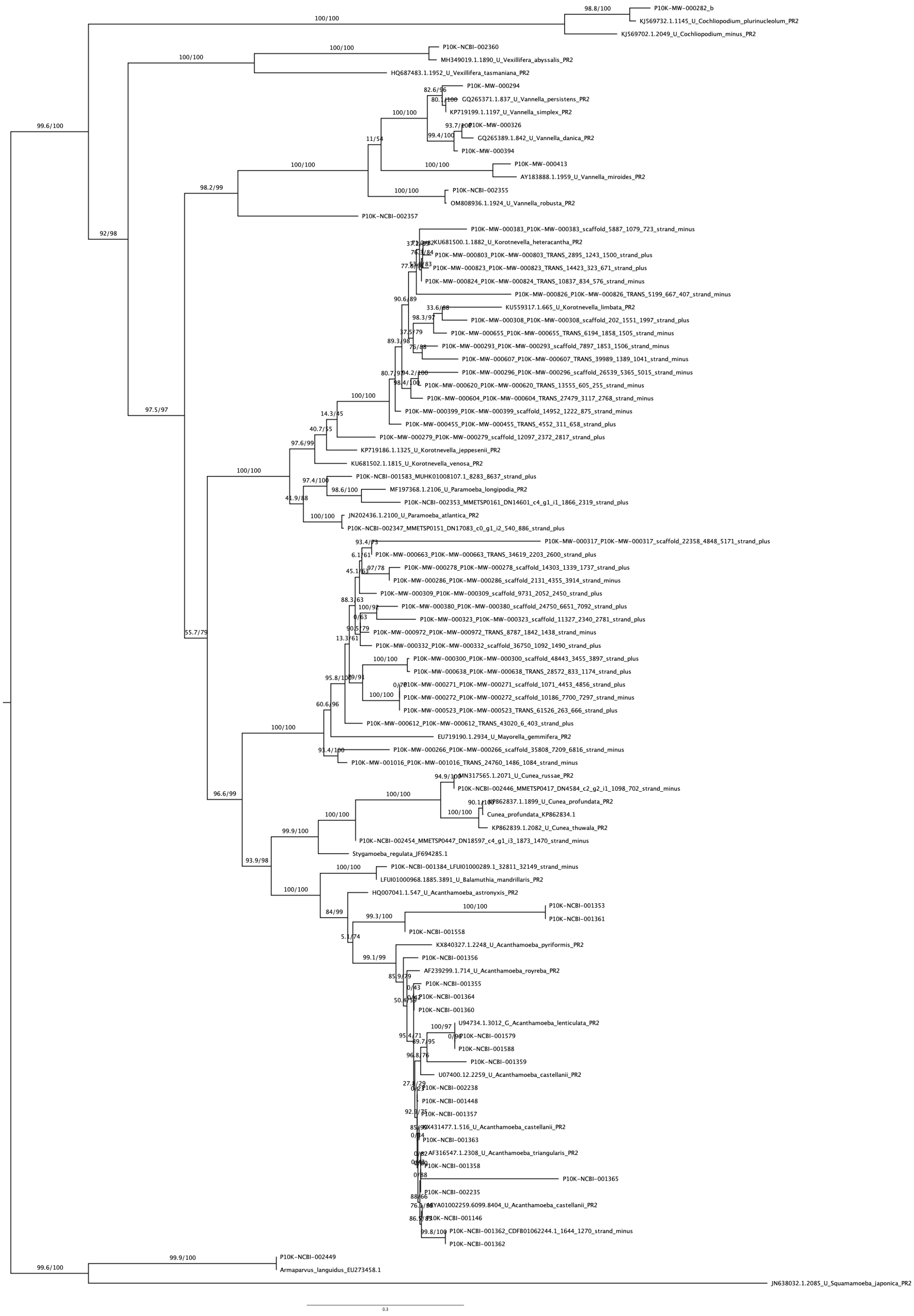
100/100
98.8/100
P10K-MW-000282_b
KJ569732.1.1145_U_Cochliopodium_plurinucleolum_PR2
KJ569702.1.2049_U_Cochliopodium_minus_PR2
100/100
P10K-NCBI-002360
MH349019.1.1890_U_Vexillifera_abyssalis_PR2
100/100
HQ687483.1.1952_U_Vexillifera_tasmaniana_PR2
99.6/100
P10K-MW-000294
82.6/96
GQ265371.1.837_U_Vannella_persistens_PR2
KP719199.1.1197_U_Vannella_simplex_PR2
100/100
P10K-MW-000326
GQ265389.1.842_U_Vannella_danica_PR2
99.4/100
P10K-MW-000394
11/54
100/100
P10K-MW-000413
100/100
AY183888.1.1959_U_Vannella_miroides_PR2
98.2/99
100/100
P10K-NCBI-002355
OM808936.1.1924_U_Vannella_robusta_PR2
92/98
P10K-NCBI-002357
P10K-MW-000383_P10K-MW-000383_scaffold_5887_1079_723_strand_minus
KU681500.1.1882_U_Korotnevella_heteracantha_PR2
P10K-MW-000803_P10K-MW-000803_TRANS_2895_1243_1500_strand_plus
P10K-MW-000823_P10K-MW-000823_TRANS_14423_323_671_strand_plus
P10K-MW-000824_P10K-MW-000824_TRANS_10837_834_576_strand_minus
P10K-MW-000826_P10K-MW-000826_TRANS_5199_667_407_strand_minus
90.6/89
33.6/88
KU559317.1.665_U_Korotnevella_limbata_PR2
98.3/97
P10K-MW-000308_P10K-MW-000308_scaffold_202_1551_1997_strand_plus
37.5/79
P10K-MW-000655_P10K-MW-000655_TRANS_6194_1858_1505_strand_minus
89.3/98
P10K-MW-000293_P10K-MW-000293_scaffold_7897_1853_1506_strand_minus
75/88
P10K-MW-000607_P10K-MW-000607_TRANS_39989_1389_1041_strand_minus
P10K-MW-000296_P10K-MW-000296_scaffold_26539_5365_5015_strand_minus
P10K-MW-000620_P10K-MW-000620_TRANS_13555_605_255_strand_minus
P10K-MW-000604_P10K-MW-000604_TRANS_27479_3117_2768_strand_minus
100/100
P10K-MW-000399_P10K-MW-000399_scaffold_14952_1222_875_strand_minus
97.5/97
P10K-MW-000455_P10K-MW-000455_TRANS_4552_311_658_strand_plus
14.3/45
40.7/55
P10K-MW-000279_P10K-MW-000279_scaffold_12097_2372_2817_strand_plus
97.6/99
KP719186.1.1325_U_Korotnevella_jeppesenii_PR2
KU681502.1.1815_U_Korotnevella_venosa_PR2
100/100
P10K-NCBI-001583_MUHK01008107.1_8283_8637_strand_plus
97.4/100
98.6/100
MF197368.1.2106_U_Paramoeba_longipodia_PR2
41.9/88
P10K-NCBI-002353_MMETSP0161_DN14601_c4_g1_i1_1866_2319_strand_plus
100/100
JN202436.1.2100_U_Paramoeba_atlantica_PR2
P10K-NCBI-002347_MMETSP0151_DN17083_c0_g1_i2_540_886_strand_plus
93.4/73
P10K-MW-000317_P10K-MW-000317_scaffold_22358_4848_5171_strand_plus
6.1/61
P10K-MW-000663_P10K-MW-000663_TRANS_34619_2203_2600_strand_plus
97/78
P10K-MW-000278_P10K-MW-000278_scaffold_14303_1339_1737_strand_plus
45.1/63
P10K-MW-000286_P10K-MW-000286_scaffold_2131_4355_3914_strand_minus
P10K-MW-000309_P10K-MW-000309_scaffold_9731_2052_2450_strand_plus
88.3/63
100/92
P10K-MW-000380_P10K-MW-000380_scaffold_24750_6651_7092_strand_plus
0/63
P10K-MW-000323_P10K-MW-000323_scaffold_11327_2340_2781_strand_plus
55.7/79
90.5/79
P10K-MW-000972_P10K-MW-000972_TRANS_8787_1842_1438_strand_minus
13.3/61
P10K-MW-000332_P10K-MW-000332_scaffold_36750_1092_1490_strand_plus
100/100
P10K-MW-000300_P10K-MW-000300_scaffold_48443_3455_3897_strand_plus
P10K-MW-000638_P10K-MW-000638_TRANS_28572_833_1174_strand_plus
P10K-MW-000271_P10K-MW-000271_scaffold_1071_4453_4856_strand_plus
100/100
P10K-MW-000272_P10K-MW-000272_scaffold_10186_7700_7297_strand_minus
P10K-MW-000523_P10K-MW-000523_TRANS_61526_263_666_strand_plus
60.6/96
P10K-MW-000612_P10K-MW-000612_TRANS_43020_6_403_strand_plus
100/100
EU719190.1.2934_U_Mayorella_gemmifera_PR2
93.4/100
P10K-MW-000266_P10K-MW-000266_scaffold_35808_7209_6816_strand_minus
P10K-MW-001016_P10K-MW-001016_TRANS_24760_1486_1084_strand_minus
MN317565.1.2071_U_Cunea_russae_PR2
96.6/99
P10K-NCBI-002446_MMETSP0417_DN4584_c2_g2_i1_1098_702_strand_minus
100/100
KP862837.1.1899_U_Cunea_profundata_PR2
100/100
Cunea_profundata_KP862834.1
100/100
KP862839.1.2082_U_Cunea_thuwala_PR2
99.9/100
P10K-NCBI-002454_MMETSP0447_DN18597_c4_g1_i3_1873_1470_strand_minus
Stygamoeba_regulata_JF694285.1
93.9/98
100/100
P10K-NCBI-001384_LFUI01000289.1_32811_32149_strand_minus
LFUI01000968.1885.3891_U_Balamuthia_mandrillaris_PR2
100/100
HQ007041.1.547_U_Acanthamoeba_astronyxis_PR2
100/100
P10K-NCBI-001353
84/99
99.3/100
P10K-NCBI-001361
P10K-NCBI-001558
5.1/74
KX840327.1.2248_U_Acanthamoeba_pyriformis_PR2
99.1/99
P10K-NCBI-001356
AF239299.1.714_U_Acanthamoeba_royreba_PR2
P10K-NCBI-001355
P10K-NCBI-001364
P10K-NCBI-001360
100/97
U94734.1.3012_G_Acanthamoeba_lenticulata_PR2
P10K-NCBI-001579
P10K-NCBI-001588
P10K-NCBI-001359
U07400.12.2259_U_Acanthamoeba_castellanii_PR2
P10K-NCBI-002238
P10K-NCBI-001448
P10K-NCBI-001357
KX431477.1.516_U_Acanthamoeba_castellanii_PR2
P10K-NCBI-001363
AF316547.1.2308_U_Acanthamoeba_triangularis_PR2
P10K-NCBI-001358
P10K-NCBI-001365
P10K-NCBI-002235
AEYA01002259.6099.8404_U_Acanthamoeba_castellanii_PR2
P10K-NCBI-001146
P10K-NCBI-001362_CDFB01062244.1_1644_1270_strand_minus
99.8/100
P10K-NCBI-001362
99.9/100
P10K-NCBI-002449
99.6/100
Armaparvus_languidus_EU273458.1
JN638032.1.2085_U_Squamamoeba_japonica_PR2
0.3

**Figure S5**. The maximum-likelihood phylogenetic tree of Small Subunit ribosomal RNA (SSU) inferred from a subset of the curated dataset presented in Figure S3, focuses on the major Amoebozoa group Discosea and includes Cutosea, a long-branch lineage that consistently falls outside Evosea in SSU rRNA phylogenies (Schuler and Brown 2019), and is shown as Figure 2B in the main text of the manuscript. Phylogenetic reconstruction was conducted using IQ-TREE v2.3.6, with ModelFinder identifying the best-fit substitution model (GTR+F+G4). Node support was assessed using both ultrafast bootstrap (UFBoot) and the Shimodaira–Hasegawa approximate likelihood ratio test (SH-aLRT). Support values are reported as SH-aLRT / UFBoot, with values ≥80/95 considered indicative of strong support.

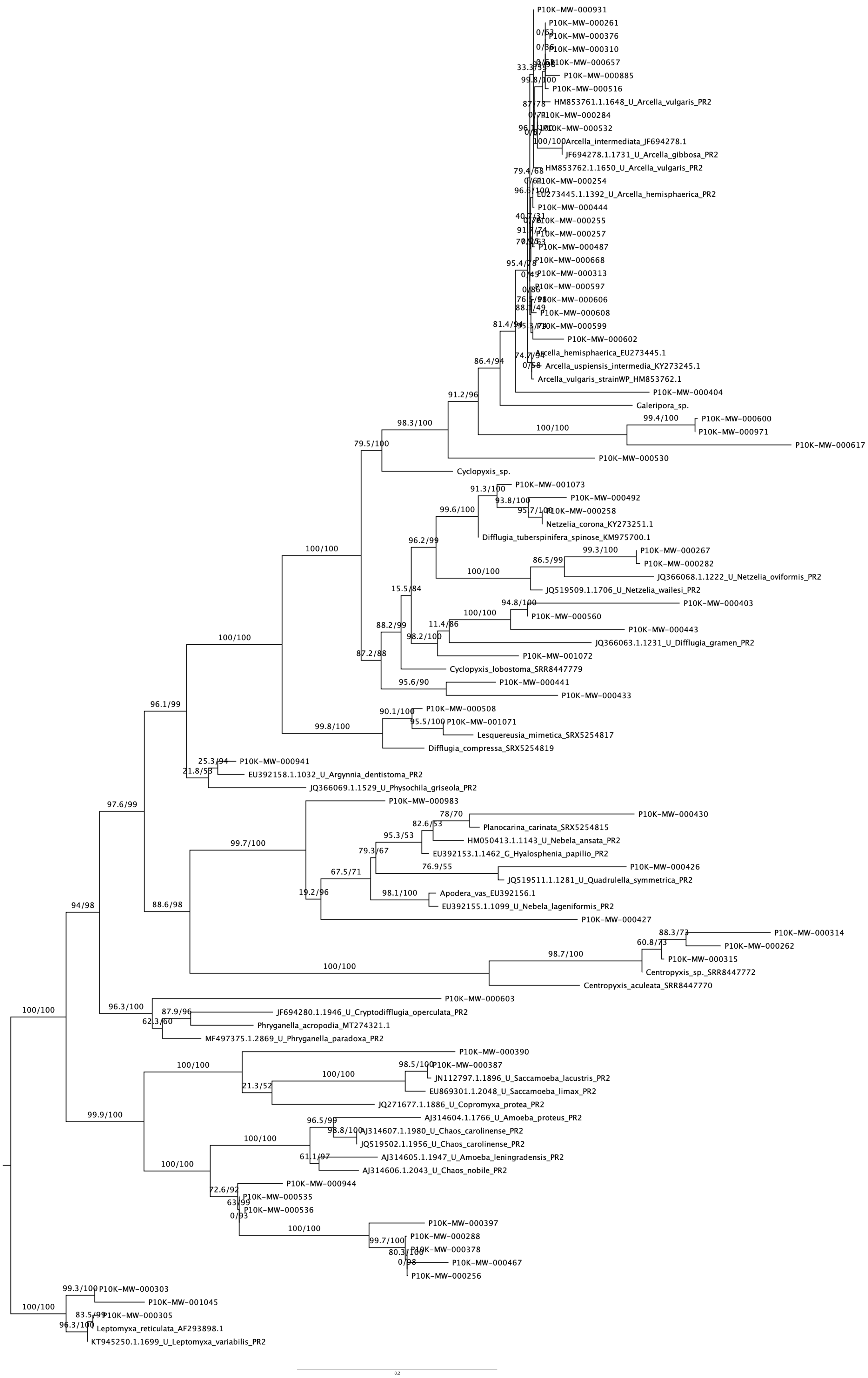
P10K-MW-000931
P10K-MW-000261
P10K-MW-000376
P10K-MW-000310
P10K-MW-000657
P10K-MW-000885
P10K-MW-000516
HM853761.1.1648_U_Arcella_vulgaris_PR2
P10K-MW-000284
P10K-MW-000532
100/100
Arcella_intermediata_JF694278.1
JF694278.1.1731_U_Arcella_gibbosa_PR2
HM853762.1.1650_U_Arcella_vulgaris_PR2
79.4/68
P10K-MW-000254
EU273445.1.1392_U_Arcella_hemisphaerica_PR2
P10K-MW-000444
P10K-MW-000255
P10K-MW-000257
P10K-MW-000487
95.4/78
P10K-MW-000668
P10K-MW-000313
P10K-MW-000597
P10K-MW-000606
P10K-MW-000608
P10K-MW-000599
P10K-MW-000602
81.4/94
Arcella_hemisphaerica_EU273445.1
74.7/94
Arcella_uspiensis_intermedia_KY273245.1
Arcella_vulgaris_strainWP_HM853762.1
86.4/94
P10K-MW-000404
Galeripora_sp.
91.2/96
100/100
99.4/100
P10K-MW-000600
P10K-MW-000971
P10K-MW-000617
98.3/100
79.5/100
P10K-MW-000530
Cyclopyxis_sp.
91.3/100
P10K-MW-001073
93.8/100
P10K-MW-000492
99.6/100
95.7/100
P10K-MW-000258
Netzelia_corona_KY273251.1
Difflugia_tuberspinifera_spinose_KM975700.1
96.2/99
99.3/100
P10K-MW-000267
86.5/99
P10K-MW-000282
100/100
JQ366068.1.1222_U_Netzelia_oviformis_PR2
JQ519509.1.1706_U_Netzelia_wailesi_PR2
15.5/84
94.8/100
P10K-MW-000403
100/100
P10K-MW-000560
11.4/86
P10K-MW-000443
88.2/99
98.2/100
JQ366063.1.1231_U_Difflugia_gramen_PR2
87.2/88
P10K-MW-001072
Cyclopyxis_lobostoma_SRR8447779
95.6/90
P10K-MW-000441
P10K-MW-000433
100/100
90.1/100
P10K-MW-000508
95.5/100
P10K-MW-001071
100/100
99.8/100
Lesquereusia_mimetica_SRX5254817
Difflugia_compressa_SRX5254819
96.1/99
25.3/94
P10K-MW-000941
21.8/53
EU392158.1.1032_U_Argynnia_dentistoma_PR2
JQ366069.1.1529_U_Physochila_griseola_PR2
97.6/99
P10K-MW-000983
78/70
P10K-MW-000430
82.6/53
Planocarina_carinata_SRX5254815
95.3/53
HM050413.1.1143_U_Nebela_ansata_PR2
99.7/100
79.3/67
EU392153.1.1462_G_Hyalosphenia_papilio_PR2
67.5/71
76.9/55
P10K-MW-000426
JQ519511.1.1281_U_Quadrulella_symmetrica_PR2
19.2/96
98.1/100
Apodera_vas_EU392156.1
EU392155.1.1099_U_Nebela_lageniformis_PR2
88.6/98
P10K-MW-000427
94/98
88.3/73
P10K-MW-000314
60.8/73
P10K-MW-000262
98.7/100
P10K-MW-000315
100/100
Centropyxis_sp._SRR8447772
Centropyxis_aculeata_SRR8447770
100/100
96.3/100
P10K-MW-000603
87.9/96
JF694280.1.1946_U_Cryptodifflugia_operculata_PR2
62.3/60
Phryganella_acropodia_MT274321.1
MF497375.1.2869_U_Phryganella_paradoxa_PR2
100/100
P10K-MW-000390
98.5/100
P10K-MW-000387
100/100
JN112797.1.1896_U_Saccamoeba_lacustris_PR2
21.3/52
EU869301.1.2048_U_Saccamoeba_limax_PR2
99.9/100
JQ271677.1.1886_U_Copromyxa_protea_PR2
96.5/99
AJ314604.1.1766_U_Amoeba_proteus_PR2
98.8/100
AJ314607.1.1980_U_Chaos_carolinense_PR2
100/100
JQ519502.1.1956_U_Chaos_carolinense_PR2
61.1/97
AJ314605.1.1947_U_Amoeba_leningradensis_PR2
100/100
AJ314606.1.2043_U_Chaos_nobile_PR2
P10K-MW-000944
72.6/92
P10K-MW-000535
63/99
P10K-MW-000536
0/93
P10K-MW-000397
100/100
P10K-MW-000288
99.7/100
P10K-MW-000378
80.3/100
P10K-MW-000467
0/98
P10K-MW-000256
99.3/100
P10K-MW-000303
100/100
P10K-MW-001045
83.5/99
P10K-MW-000305
96.3/100
Leptomyxa_reticulata_AF293898.1
KT945250.1.1699_U_Leptomyxa_variabilis_PR2
0.2

**Figure S6**. The maximum-likelihood phylogenetic tree of Small Subunit ribosomal RNA (SSU) inferred from a subset of the curated dataset presented in Figure S3, focuses on the major Amoebozoa group Tubulinea and is shown as Figure 3A in the main text of the manuscript. Phylogenetic reconstruction was conducted using IQ-TREE v2.3.6, with ModelFinder identifying the best-fit substitution model (TIM3e+G4). Node support was assessed using both ultrafast bootstrap (UFBoot) and the Shimodaira–Hasegawa approximate likelihood ratio test (SH-aLRT). Support values are reported as SH-aLRT / UFBoot, with values ≥80/95 considered indicative of strong support.

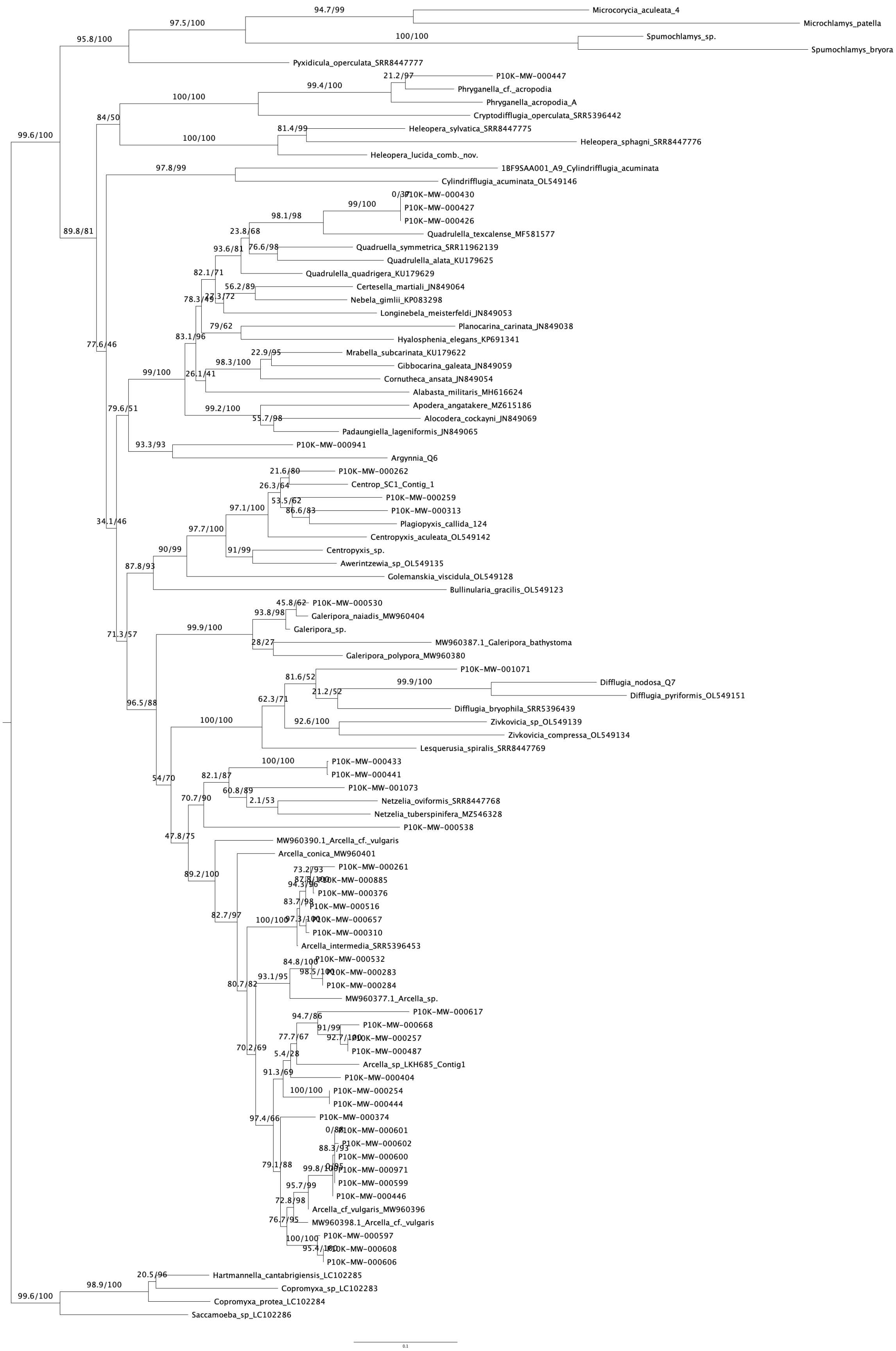

Microcorycia_aculeata_4
Microchlamys_patella
Spumochlamys_sp.
Spumochlamys_bryora
Pyxidicula_operculata_SRR8447777
P10K-MW-000447
Phryganella_cf._acropodia
Phryganella_acropodia_A
Cryptodifflugia_operculata_SRR5396442
Heleopera_sylvatica_SRR8447775
Heleopera_sphagni_SRR8447776
Heleopera_lucida_comb._nov.
1BF9SAA001_A9_Cylindrifflugia_acuminata
Cylindrifflugia_acuminata_OL549146
P10K-MW-000430
P10K-MW-000427
P10K-MW-000426
Quadrulella_texcalense_MF581577
Quadruella_symmetrica_SRR11962139
Quadrulella_alata_KU179625
Quadrulella_quadrigera_KU179629
Certesella_martiali_JN849064
Nebela_gimlii_KP083298
Longinebela_meisterfeldi_JN849053
Planocarina_carinata_JN849038
Hyalosphenia_elegans_KP691341
Mrabella_subcarinata_KU179622
Gibbocarina_galeata_JN849059
Cornutheca_ansata_JN849054
Alabasta_militaris_MH616624
Apodera_angatakere_MZ615186
Alocodera_cockayni_JN849069
Padaungiella_lageniformis_JN849065
P10K-MW-000941
Argynnia_Q6
P10K-MW-000262
Centrop_SC1_Contig_1
P10K-MW-000259
P10K-MW-000313
Plagiopyxis_callida_124
Centropyxis_aculeata_OL549142
Centropyxis_sp.
Awerintzewia_sp_OL549135
Golemanskia_viscidula_OL549128
Bullinularia_gracilis_OL549123
P10K-MW-000530
Galeripora_naiadis_MW960404
Galeripora_sp.
MW960387.1_Galeripora_bathystoma
Galeripora_polypora_MW960380
P10K-MW-001071
Difflugia_nodosa_Q7
Difflugia_pyriformis_OL549151
Difflugia_bryophila_SRR5396439
Zivkovicia_sp_OL549139
Zivkovicia_compressa_OL549134
Lesquerusia_spiralis_SRR8447769
P10K-MW-000433
P10K-MW-000441
P10K-MW-001073
Netzelia_oviformis_SRR8447768
Netzelia_tuberspinifera_MZ546328
P10K-MW-000538
MW960390.1_Arcella_cf._vulgaris
Arcella_conica_MW960401
P10K-MW-000261
P10K-MW-000885
P10K-MW-000376
P10K-MW-000516
P10K-MW-000657
P10K-MW-000310
Arcella_intermedia_SRR5396453
P10K-MW-000532
P10K-MW-000283
P10K-MW-000284
MW960377.1_Arcella_sp.
P10K-MW-000617
P10K-MW-000668
P10K-MW-000257
P10K-MW-000487
Arcella_sp_LKH685_Contig1
P10K-MW-000404
P10K-MW-000254
P10K-MW-000444
P10K-MW-000374
P10K-MW-000601
P10K-MW-000602
P10K-MW-000600
P10K-MW-000971
P10K-MW-000599
P10K-MW-000446
Arcella_cf_vulgaris_MW960396
MW960398.1_Arcella_cf._vulgaris
P10K-MW-000597
P10K-MW-000608
P10K-MW-000606
Hartmannella_cantabrigiensis_LC102285
Copromyxa_sp_LC102283
Copromyxa_protea_LC102284
Saccamoeba_sp_LC102286
0.1

**Figure S7.** The maximum-likelihood phylogenetic tree of cytochrome c oxidase subunit I (COI) inferred from a curated dataset generated in the present study, focusing on Arcellinida order (Tubulinea:Amoebozoa) comprising COI sequences retrieved from genomes and transcriptomes available in the P10K database, along with reference sequences made available by previous studies and is shown as Figure 3B in the main text of the manuscript. Phylogenetic reconstruction was conducted using IQ-TREE v2.3.6, with ModelFinder identifying the best-fit substitution model (GTR+F+I+G4). Node support was assessed using both ultrafast bootstrap (UFBoot) and the Shimodaira–Hasegawa approximate likelihood ratio test (SH-aLRT). Support values are reported as SH-aLRT / UFBoot, with values ≥80/95 considered indicative of strong support.

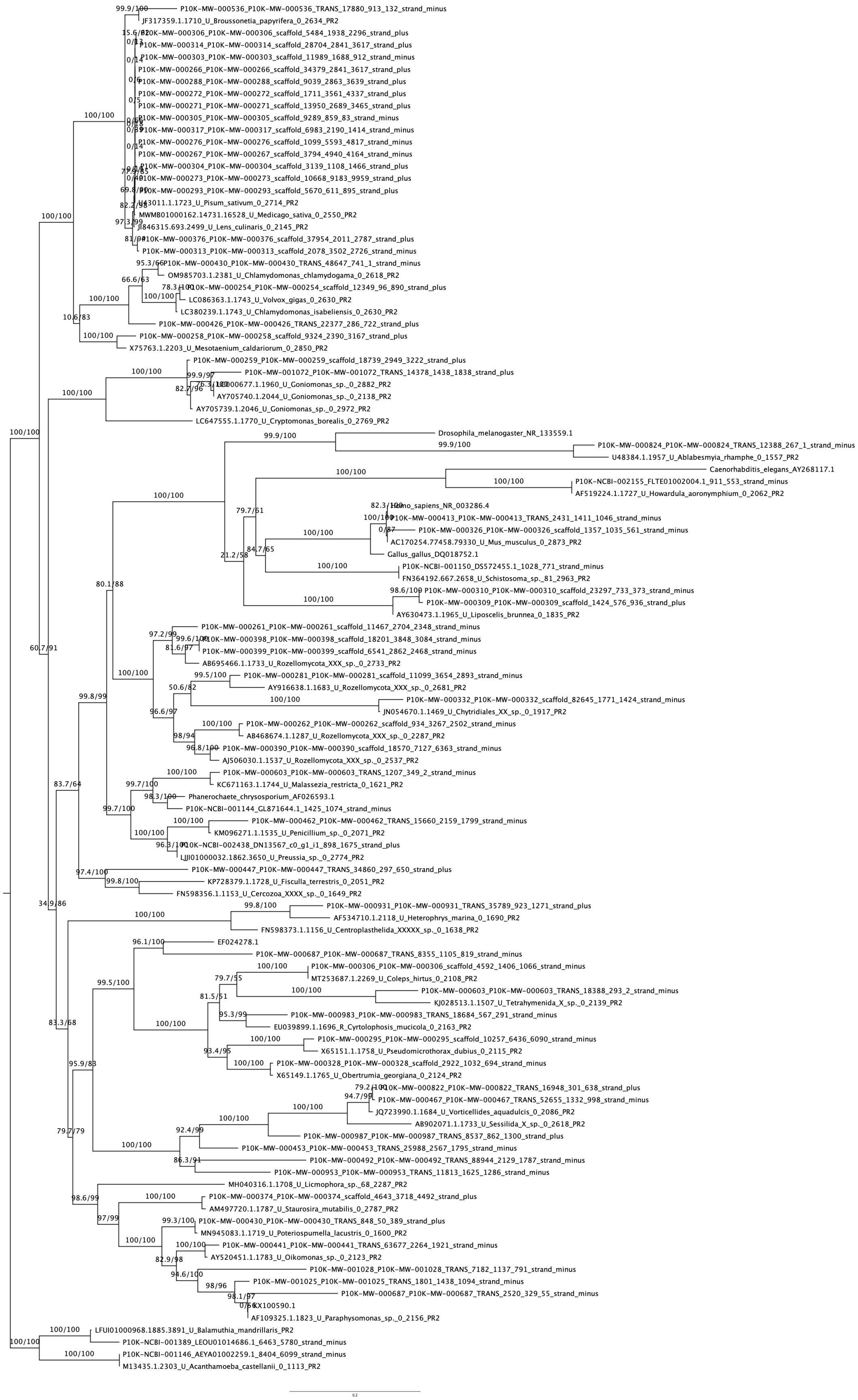
P10K-MW-000536_P10K-MW-000536_TRANS_17880_913_132_strand_minus
JF317359.1.1710_U_Broussonetia_papyrifera_0_2634_PR2
P10K-MW-000306_P10K-MW-000306_scaffold_5484_1938_2296_strand_plus
P10K-MW-000314_P10K-MW-000314_scaffold_28704_2841_3617_strand_plus
P10K-MW-000303_P10K-MW-000303_scaffold_11989_1688_912_strand_minus
P10K-MW-000266_P10K-MW-000266_scaffold_34379_2841_3617_strand_plus
P10K-MW-000288_P10K-MW-000288_scaffold_9039_2863_3639_strand_plus
P10K-MW-000272_P10K-MW-000272_scaffold_1711_3561_4337_strand_plus
P10K-MW-000271_P10K-MW-000271_scaffold_13950_2689_3465_strand_plus
P10K-MW-000305_P10K-MW-000305_scaffold_9289_859_83_strand_minus
P10K-MW-000317_P10K-MW-000317_scaffold_6983_2190_1414_strand_minus
P10K-MW-000276_P10K-MW-000276_scaffold_1099_5593_4817_strand_minus
P10K-MW-000267_P10K-MW-000267_scaffold_3794_4940_4164_strand_minus
P10K-MW-000304_P10K-MW-000304_scaffold_3139_1108_1466_strand_plus
P10K-MW-000273_P10K-MW-000273_scaffold_10668_9183_9959_strand_plus
P10K-MW-000293_P10K-MW-000293_scaffold_5670_611_895_strand_plus
U43011.1.1723_U_Pisum_sativum_0_2714_PR2
MWMB01000162.14731.16528_U_Medicago_sativa_0_2550_PR2
JJ846315.693.2499_U_Lens_culinaris_0_2145_PR2
P10K-MW-000376_P10K-MW-000376_scaffold_37954_2011_2787_strand_plus
P10K-MW-000313_P10K-MW-000313_scaffold_2078_3502_2726_strand_minus
P10K-MW-000430_P10K-MW-000430_TRANS_48647_741_1_strand_minus
OM985703.1.2381_U_Chlamydomonas_chlamydogama_0_2618_PR2
P10K-MW-000254_P10K-MW-000254_scaffold_12349_96_890_strand_plus
LC086363.1.1743_U_Volvox_gigas_0_2630_PR2
LC380239.1.1743_U_Chlamydomonas_isabeliensis_0_2630_PR2
P10K-MW-000426_P10K-MW-000426_TRANS_22377_286_722_strand_plus
P10K-MW-000258_P10K-MW-000258_scaffold_9324_2390_3167_strand_plus
X75763.1.2203_U_Mesotaenium_caldariorum_0_2850_PR2
P10K-MW-000259_P10K-MW-000259_scaffold_18739_2949_3222_strand_plus
P10K-MW-001072_P10K-MW-001072_TRANS_14378_1438_1838_strand_plus
AY705740.1.2044_U_Goniomonas_sp._0_2138_PR2
AY705739.1.2046_U_Goniomonas_sp._0_2972_PR2
LC647555.1.1770_U_Cryptomonas_borealis_0_2769_PR2
Drosophila_melanogaster_NR_133559.1
P10K-MW-000824_P10K-MW-000824_TRANS_12388_267_1_strand_minus
U48384.1.1957_U_Ablabesmyia_rhamphe_0_1557_PR2
Caenorhabditis_elegans_AY268117.1
P10K-NCBI-002155_FLTE01002004.1_911_553_strand_minus
AF519224.1.1727_U_Howardula_aoronymphium_0_2062_PR2
P10K-MW-000413_P10K-MW-000413_TRANS_2431_1411_1046_strand_minus
P10K-MW-000326_P10K-MW-000326_scaffold_1357_1035_561_strand_minus
AC170254.77458.79330_U_Mus_musculus_0_2873_PR2
Gallus_gallus_DQ018752.1
P10K-NCBI-001150_DS572455.1_1028_771_strand_minus
FN364192.667.2658_U_Schistosoma_sp._81_2963_PR2
P10K-MW-000310_P10K-MW-000310_scaffold_23297_733_373_strand_minus
P10K-MW-000309_P10K-MW-000309_scaffold_1424_576_936_strand_plus
AY630473.1.1965_U_Liposcelis_brunnea_0_1835_PR2
P10K-MW-000261_P10K-MW-000261_scaffold_11467_2704_2348_strand_minus
P10K-MW-000398_P10K-MW-000398_scaffold_18201_3848_3084_strand_minus
P10K-MW-000399_P10K-MW-000399_scaffold_6541_2862_2468_strand_minus
AB695466.1.1733_U_Rozellomycota_XXX_sp._0_2733_PR2
P10K-MW-000281_P10K-MW-000281_scaffold_11099_3654_2893_strand_minus
AY916638.1.1683_U_Rozellomycota_XXX_sp._0_2681_PR2
P10K-MW-000332_P10K-MW-000332_scaffold_82645_1771_1424_strand_minus
JN054670.1.1469_U_Chytridiales_XX_sp._0_1917_PR2
P10K-MW-000262_P10K-MW-000262_scaffold_934_3267_2502_strand_minus
AB468674.1.1287_U_Rozellomycota_XXX_sp._0_2287_PR2
P10K-MW-000390_P10K-MW-000390_scaffold_18570_7127_6363_strand_minus
AJ506030.1.1537_U_Rozellomycota_XXX_sp._0_2537_PR2
P10K-MW-000603_P10K-MW-000603_TRANS_1207_349_2_strand_minus
KC671163.1.1744_U_Malassezia_restricta_0_1621_PR2
Phanerochaete_chrysosporium_AF026593.1
P10K-NCBI-001144_GL871644.1_1425_1074_strand_minus
P10K-MW-000462_P10K-MW-000462_TRANS_15660_2159_1799_strand_minus
KM096271.1.1535_U_Penicillium_sp._0_2071_PR2
P10K-NCBI-002438_DN13567_c0_g1_i1_898_1675_strand_plus
LJJI01000032.1862.3650_U_Preussia_sp._0_2774_PR2
P10K-MW-000447_P10K-MW-000447_TRANS_34860_297_650_strand_plus
KP728379.1.1728_U_Fisculla_terrestris_0_2051_PR2
FN598356.1.1153_U_Cercozoa_XXXX_sp._0_1649_PR2
P10K-MW-000931_P10K-MW-000931_TRANS_35789_923_1271_strand_plus
AF534710.1.2118_U_Heterophrys_marina_0_1690_PR2
FN598373.1.1156_U_Centroplasthelida_XXXXX_sp._0_1638_PR2
EF024278.1
P10K-MW-000687_P10K-MW-000687_TRANS_8355_1105_819_strand_minus
P10K-MW-000306_P10K-MW-000306_scaffold_4592_1406_1066_strand_minus
MT253687.1.2269_U_Coleps_hirtus_0_2108_PR2
P10K-MW-000603_P10K-MW-000603_TRANS_18388_293_2_strand_minus
KJ028513.1.1507_U_Tetrahymenida_X_sp._0_2139_PR2
P10K-MW-000983_P10K-MW-000983_TRANS_18684_567_291_strand_minus
EU039899.1.1696_R_Cyrtolophosis_mucicola_0_2163_PR2
P10K-MW-000295_P10K-MW-000295_scaffold_10257_6436_6090_strand_minus
X65151.1.1758_U_Pseudomicrothorax_dubius_0_2115_PR2
P10K-MW-000328_P10K-MW-000328_scaffold_2922_1032_694_strand_minus
X65149.1.1765_U_Obertrumia_georgiana_0_2124_PR2
P10K-MW-000822_P10K-MW-000822_TRANS_16948_301_638_strand_plus
P10K-MW-000467_P10K-MW-000467_TRANS_52655_1332_998_strand_minus
JQ723990.1.1684_U_Vorticellides_aquadulcis_0_2086_PR2
AB902071.1.1733_U_Sessilida_X_sp._0_2618_PR2
P10K-MW-000987_P10K-MW-000987_TRANS_8537_862_1300_strand_plus
P10K-MW-000453_P10K-MW-000453_TRANS_25988_2567_1795_strand_minus
P10K-MW-000492_P10K-MW-000492_TRANS_88944_2129_1787_strand_minus
P10K-MW-000953_P10K-MW-000953_TRANS_11813_1625_1286_strand_minus
MH040316.1.1708_U_Licmophora_sp._68_2287_PR2
P10K-MW-000374_P10K-MW-000374_scaffold_4643_3718_4492_strand_plus
AM497720.1.1787_U_Staurosira_mutabilis_0_2787_PR2
P10K-MW-000430_P10K-MW-000430_TRANS_848_50_389_strand_plus
MN945083.1.1719_U_Poteriospumella_lacustris_0_1600_PR2
P10K-MW-000441_P10K-MW-000441_TRANS_63677_2264_1921_strand_minus
AY520451.1.1783_U_Oikomonas_sp._0_2123_PR2
P10K-MW-001028_P10K-MW-001028_TRANS_7182_1137_791_strand_minus
P10K-MW-001025_P10K-MW-001025_TRANS_1801_1438_1094_strand_minus
P10K-MW-000687_P10K-MW-000687_TRANS_2520_329_55_strand_minus
KX100590.1
AF109325.1.1823_U_Paraphysomonas_sp._0_2156_PR2
LFUI01000968.1885.3891_U_Balamuthia_mandrillaris_PR2
P10K-NCBI-001389_LEOU01014686.1_6463_5780_strand_minus
P10K-NCBI-001146_AEYA01002259.1_8404_6099_strand_minus
M13435.1.2303_U_Acanthamoeba_castellanii_0_1113_PR2
0.2

**Figure S8.** The maximum-likelihood phylogenetic tree of Small Subunit ribosomal RNA (SSU), inferred from a curated dataset generated in the present study, comprises SSU sequences of putative non-Amoebozoan contaminants retrieved from genomes and transcriptomes available in the P10K database, along with reference sequences from the PR$^2$ database (indicated by PR$^2$ IDs), identified through BLAST+ similarity searches using the putative contaminant SSU sequences as queries. This tree is presented as Figure 4 in the main text of the manuscript. Phylogenetic reconstruction was conducted using IQ-TREE v2.3.6, with ModelFinder identifying the best-fit substitution model (TN+F+I+G4). Node support was assessed using both ultrafast bootstrap (UFBoot) and the Shimodaira–Hasegawa approximate likelihood ratio test (SH-aLRT). Support values are reported as SH-aLRT / UFBoot, with values ≥80/95 considered indicative of strong support.

### Supplementary dataset legends

**Data S1**. Representative output from running the CSI-SSU tool on the 2,960 assemblies from the P10K dataset, encompassing all supergroups considered in this study. The data include the parsed_blast output directory, containing parsed and filtered SSU sequences, the vsearch directory, containing summaries of chimeric search results, and the summary directory, containing final summary reports and phylogenetic trees with sequence placements (PDF format).

**Data S2. The master SSU rDNA/rRNA dataset constructed in this study (see *Methods* section from the main text).** This dataset, prior to curation (e.g., removal of duplicate and short sequences based on a preliminary phylogenetic reconstruction), includes all SSU sequences identified and retrieved from Amoebozoan genomic data available in the P10K database (indicated by P10K sample IDs), along with SSU rDNA/rRNA sequences from previous studies and the PR$^2$ database (indicated by PR$^2$ IDs) (Kang et al. 2017; Ribeiro et al. 2023; Porfirio-Sousa et al. 2024). It also incorporates reference sequences of non-amoebozoan taxa from the PR$^2$ database used for contamination screening of P10K genomic data.

**Data S3. The curated SSU rDNA/rRNA master Amoebozoa dataset focusing on the major amoebozoan groups Tubulinea, Evosea, and Discosea.** This is the dataset used to construct the phylogenetic tree presented in the **Figure S3** and the datasets **Data S4 – S6**.

**Data S4. The curated subset of the SSU rDNA/rRNA master dataset focusing on the major Amoebozoa group Evosea.** This is the dataset used to construct the phylogenetic tree presented in **Figure 2A** and **Figure S4**.

**Data S5. The curated subset of the SSU rDNA/rRNA master dataset focusing on the major Amoebozoa group Discosea.** This is the dataset used to construct the phylogenetic tree presented in **Figure 2B** and **Figure S5**.

**Data S6. The curated subset of the SSU rDNA/rRNA master dataset focusing on the major Amoebozoa group Tubulinea.** This is the dataset used to construct the phylogenetic tree presented in **Figure 3A** and **Figure S6**.

**Data S7. The curated cytochrome c oxidase subunit I (COI) dataset focusing on the order Arcellinida (Tubulinea:Amoebozoa).** This is the dataset used to construct the phylogenetic tree presented in **Figure 3B** and **Figure S7**.

**Data S8. The curated subset of the SSU rDNA/rRNA master dataset focusing on the putative non-amoebozoan contaminants.** This is the contamination screening dataset used to construct the phylogenetic tree presented in **Figure 4** and **Figure S8**.

### References


Gao Xinxin, Chen K., Xiong J., Zou D., Yang F., Ma Y., Jiang C., Gao Xiaoxuan, Wang G., Gu S., Zhang P., Luo S., Huang K., Bao Y., Zhang Z., Ma L. & Miao W. 2024. The P10K database: a data portal for the protist 10 000 genomes project. *Nucleic Acids Research*, **52**:D747–D755.

Kang S., Tice A. K., Spiegel F. W., Silberman J. D., Pánek T., Čepička I., Kostka M., Kosakyan A., Alcântara D. M. C., Roger A. J., Shadwick L. L., Smirnov A., Kudryavtsev A., Lahr D. J. G. & Brown M. W. 2017. Between a Pod and a Hard Test: The Deep Evolution of Amoebae. *Molecular Biology and Evolution*, **34**:2258–2270.

Porfirio-Sousa A. L., Tice A. K., Morais L., Ribeiro G. M., Blandenier Q., Dumack K., Eglit Y., Fry N. W., Gomes E Souza M. B., Henderson T. C., Kleitz-Singleton F., Singer D., Brown M. W. & Lahr D. J. G. 2024. Amoebozoan testate amoebae illuminate the diversity of heterotrophs and the complexity of ecosystems throughout geological time. *Proceedings of the National Academy of Sciences*, **121**:e2319628121.

Ribeiro G. M., Useros F., Dumack K., González-Miguéns R., Siemensma F., Porfírio-Sousa A. L., Soler-Zamora C., Pedro Barbosa Alcino J., Lahr D. J. G. & Lara E. 2023. Expansion of the cytochrome C oxidase subunit I database and description of four new lobose testate amoebae species (Amoebozoa; Arcellinida). *European Journal of Protistology*, **91**:126013.
